# GATA4 loss promotes mutant *Kras*-driven pancreatic ductal adenocarcinoma in the absence of canonical precursor lesions

**DOI:** 10.64898/2026.05.22.724502

**Authors:** Francesc Madriles, Monica P. de Andres, Mikhail Chesnokov, Jaime Martinez de Villarreal, Direna Alonso-Curbelo, Natalia del Pozo, Mar Iglesias, Enrique Carrillo-de-Santa-Pau, Juan Iovanna, Francisco Soriano, Ana Cuadrado, Miriam Marques, Javier Muñoz, Irene Esposito, Paola Martinelli, Scott W. Lowe, Francisco X. Real

## Abstract

GATA6 and GATA4 play key roles in pancreatic development and are essential to maintain the classical transcriptional program in pancreatic ductal adenocarcinoma (PDAC). Using genetic mouse models we show that, in contrast to GATA6, GATA4 is dispensable for the maintenance of acinar homeostasis in the adult pancreas. Deletion of *Gata4* in mice expressing mutant *Kras* in the embryonic pancreas (KG4C) leads to PDAC development in the absence of tissue remodeling, pancreatic intraepithelial neoplasia (PanIN), or other canonical precursor lesions present in *Gata4*-proficient (KC) mice. Similar observations were made when *Gata4* was selectively inactivated in adult, *Kras*-mutant, acinar cells. We identify Pale Acinar Lesions (PALes) as a previously unrecognized pancreatic lesion, distinct from acino-ductal metaplasia (ADM) and PanINs, present in KC and KG4C mice but not in wild type mice. PALes display weak expression of acinar and ductal markers and lack mucins; they have lower proliferation rates than PanINs. RNA-seq and ChIP-seq reveal that GATA4 and GATA6 partially share genomic binding sites and transcriptomic effects, but they exert opposing influences on mutant *Kras*–induced, haematopoietic cell–dependent, transcriptional inflammatory programs. Adenoviral-mediated pancreatic expression of IL17 restored the formation of ductal lesions in KG4C mice but failed to rescue PanIN development. Our data indicate that GATA4 functions through the coordinated action of multiple inflammatory factors that are required for ADM/PanIN formation but are dispensable for PDAC development. Collectively, these findings challenge current paradigms of PDAC initiation and progression.

## INTRODUCTION

Pancreatic ductal adenocarcinoma (PDAC) is a deadly cancer, even when diagnosed at early stages. Patients with PDAC have a 5-year survival rate of only 10% (Siegel et al., 2021) and most patients who undergo therapy with curative intent recur within 2 years due to distant metastases (Bray et al., 2024). It is, therefore, urgent to improve our understanding of how PDAC develops and evolves.

PDAC genomic analyses have revealed that: (1) more than 90% of tumors harbor activating mutations in *KRAS* and inactivating alterations in *INK4A*, *TP53*, and *SMAD4* (or other genes in their corresponding pathways), (2) most tumors contain additional “private” alterations in genes that are involved in <10% of tumors, (3) while tumors vary widely regarding copy number alterations and structural aberrations, these are pervasive in a large fraction of PDAC, (4) each of these alterations may arise gradually and independently (Bailey et al., 2016; Chan-Seng-Yue et al., 2020; Lüttges et al., 2001; Moskaluk et al., 1997; Notta et al., 2016; Waddell et al., 2015; Wilentz et al., 1998, 2000). The analysis of tumor cell-enriched genomes has unveiled that whole genome duplications, complex rearrangements, and chromothripsis events can inactivate simultaneously multiple tumor suppressors, providing support to a catastrophic model of tumor progression as a major contributor to invasion and metastases (Notta et al., 2016; Real, 2003). These diverse and complex tumor evolution patterns have recently been highlighted with single cell resolution (Zhang et al., 2026).

PDAC has been proposed to develop from Pancreatic Intraepithelial Neoplasia (PanINs), atypical flat lesions (AFLs), or mucinous cystic neoplasms including the most common intraductal papillary mucinous neoplasms (IPMNs) (reviewed in Pedro & Wood, 2025). Unlike PanINs and AFLs, which are small, IPMNs can be detected using non-invasive imaging techniques. PanINs can be flat or papillary and are classified as low- or high-grade. A hallmark of low-grade PanINs is the presence of mucins. Studies dating back several decades have shown that mucinous hyperplasia (an old name for low-grade PanINs) occurs in aged individuals without pancreatic disease (Allen-Mersh, 1985). More recent work has confirmed that low-grade PanINs are pervasive in the non-neoplastic pancreas of organ donors, including young adults. Importantly, they almost invariably contain one or more mutant *KRAS* allele and have a low risk of progressing to PDAC (Braxton et al., 2024; Carpenter et al., 2023; Kanda et al., 2012; Matsuda et al., 2017). On the other hand, high-grade PanINs are rare/exceptional in the absence of PDAC (Hosoda et al., 2017; Real, 2003). AFLs are highly proliferative tubular lesions that arise in areas of acinar-to-ductal metaplasia (ADM). They are surrounded by a loose but highly cellular stroma and show cytological atypia (von Figura et al., 2017). Both PanINs and AFLs have been described in patients with familial pancreatic cancer (Aichler et al., 2012).

Genetically engineered mouse models (GEMM) have proven extremely valuable to address key questions linking the genetic drivers and cell of origin of PDAC, the resulting preneoplastic lesions, and the mechanisms involved in tumor progression. Selective activation of mutant *Kras* in embryonic acinar cells can give rise to PanINs and AFLs whereas adult acinar cells are largely resistant to the oncogenic effects of mutant *Kras* (Guerra et al., 2007; Hingorani et al., 2003; von Figura et al., 2017). However, acute and chronic inflammation promote PDAC development (Carrière et al., 2011; Flandez et al., 2014; Guerra et al., 2007; Hingorani et al., 2003; von Figura et al., 2017). Cell-of-origin, somatic genetic alterations, and non-genetic factors contribute to shape the phenotype of tumors (Ferreira et al., 2017; Lee et al., 2019; Martinelli et al., 2017). Transcriptomic analyses of human and murine PDAC have established the existence of a range of molecular phenotypes, including a canonical cell identity program (i.e., classical), a basal-squamous program, and mesenchymal and neuronal programs (Bailey et al., 2016; Chan-Seng-Yue et al., 2020; Collisson et al., 2011; Moffitt et al., 2015; Mueller et al., 2018a; Shiau et al., 2024). For example, acinar cell-derived PDAC generally display a classical phenotype. In contrast, ductal cells appear less sensitive to mutant *Kras* and give rise predominantly to PDAC with a more basal phenotype (Ferreira et al., 2017; Lee et al., 2019). These programs are driven by specific transcription factors (e.g., GATA6 and HNFs, classical; ΔNp63 and MYC, basal/mesenchymal) (Kalisz et al., 2020; Maia-Silva et al., 2024; Martinelli et al., 2017; Mueller et al., 2018b; Somerville et al., 2018) that play an important role in tumor progression and drug sensitivity, including the new KRAS inhibitors (Dilly et al., 2024; Parassiadis & Johnsen, 2026; Singhal et al., 2024).

Using GEMMs, we and others have provided evidence on the role of genes involved in pancreatic differentiation in PDAC pathogenesis. Among them is GATA6, a key regulator of pancreatic lineage fidelity in PDAC (Kloesch et al., 2022; Martinelli et al., 2017) and we have recently shown that GATA4 cooperates with GATA6 to maintain the classical phenotype (de Andrés et al., 2023). GATA proteins are a family of master transcription factor regulators involved in organogenesis and differentiation (Tremblay et al., 2018). GATA4 and GATA6 are the main members of the family expressed in the human and murine pancreas (Villamayor et al., 2020). We and others have extensively studied the role of *Gata6* in pancreatic initiation and progression using GEMMs (Kloesch et al., 2022; Martinelli et al., 2016, 2017; Zhong et al., 2026). However, despite the high expression of GATA4 in acinar cells, its role in mutant *Kras*-driven PDAC initiation/progression has not been addressed. Here, we use conditional *Gata4* knockout mice to analyze the impact of its deletion in embryonic or adult pancreas and show that, in the presence of a mutant *Kras* allele, its inactivation leads to a distinct phenotype characterized by PDAC development in the absence of canonical PanIN precursor lesions. Bioinformatics analyses show that GATA4 is required for a mutant *Kras*-driven transcriptional inflammatory response that is critical for PanIN formation, but not for PDAC.

## RESULTS

### *Gata4* loss in the pancreas has a transcriptomic impact but is largely dispensable for exocrine homeostasis and response to damage

In normal mouse pancreas, GATA4 is expressed in acinar cells but it is undetectable in ductal and endocrine cells (**Fig. 1A**). To assess the impact of *Gata4* deletion on pancreatic epithelial cells and compare the effects with those of *Gata6* deletion, we generated *Ptf1a^+/Cre^;Gata4^lox/lox^* mice (*Gata4*^P-/-^). Extensive recombination was demonstrated by lack of GATA4 expression both at the protein and RNA levels (**Fig. 1A, Suppl. Fig. 1A**).

**Figure 1.**
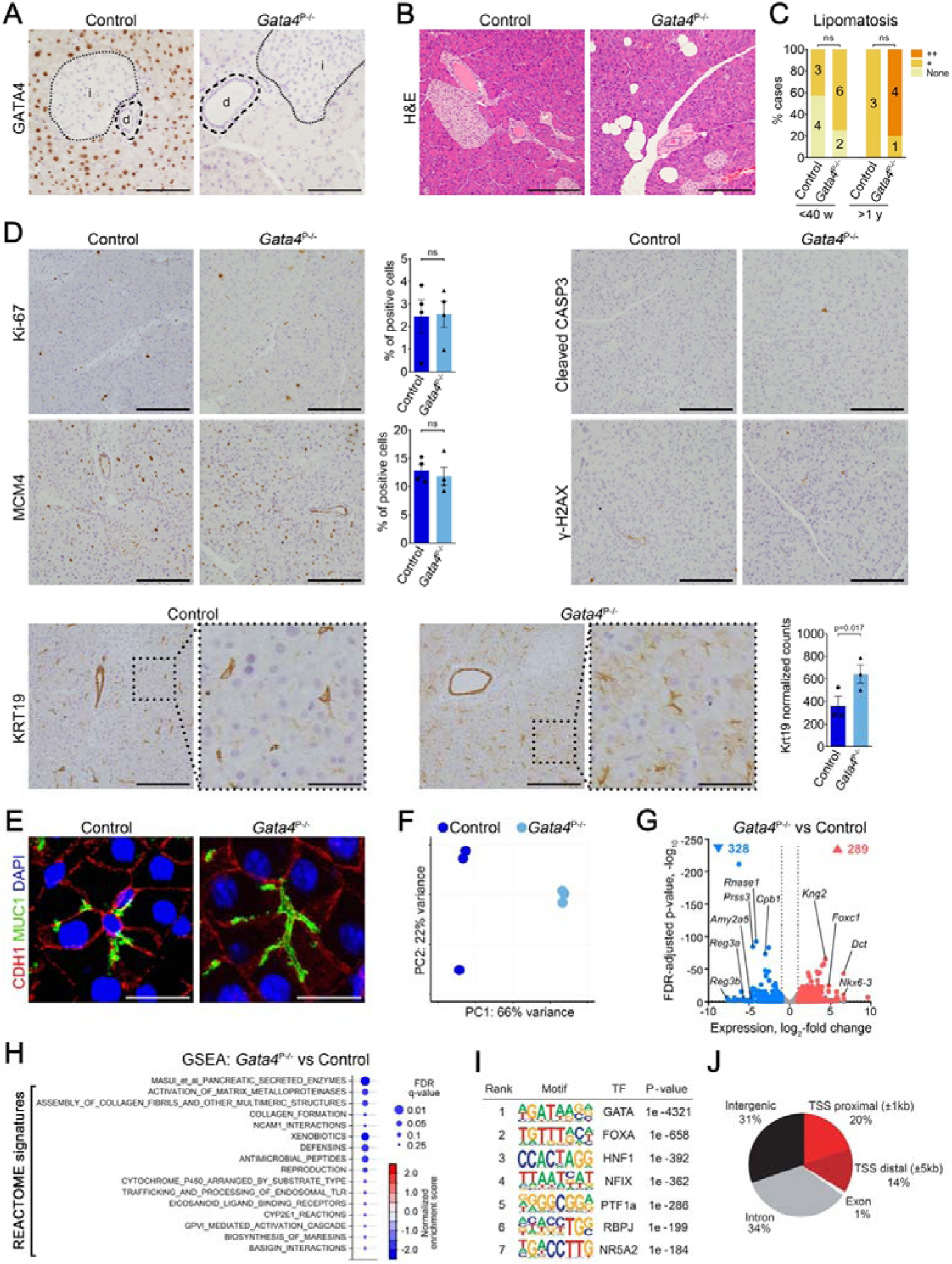
*Gata4* inactivation has a modest impact on pancreas histology and transcriptome. **(A)** GATA4 is expressed in acinar but not in islet (i) or ductal (d) cells. Upon recombination, GATA4 expression is lost in acinar cells of *Gata4*^P-/-^ mice. Scale bar, 100 µm. **(B)** H&E staining showing that the pancreas of 30-week-old *Gata4*^P-/-^ mice is histologically normal, except for a mild increase in lipomatosis and a slightly lower eosinophilia. Scale bar, 200 µm. **(C)** Quantification of lipomatosis in *Gata4*^P-/-^ pancreas showing a non-statistically significant increase both at <40 weeks (P=0.315) and >1 year (P=0.143). **(D)** IHC staining for markers of proliferation (Ki-67, MCM4), DNA damage (γ-H2AX), apoptosis (Cleaved CASP3) (all, non-significant), and ductal cells (KRT19; RNA quantification by RNA-seq). Scale bars: 200 µm (low magnification) and 50 µm (high magnification). **(E)** IF staining for cell polarity markers (E-cadherin, MUC1) shows similar expression in *Gata4*^P-/-^ and control mice. Scale bar, 20 µm. **(F)** PCA plot of RNA-seq data from control (n=3) and *Gata4*^P-/-^ (n=3) pancreata. **(G)** Volcano plot displaying differential gene expression between *Gata4*^P-/-^ and control pancreata. **(H)** Transcriptomic signatures affected in *Gata4*^P-/-^ pancreata. **(I)** Top motifs enriched in GATA4 genomic binding sites identified by ChIP-seq. **(J)** Genomic distribution of GATA4 binding sites identified by ChIP-Seq.

The pancreas of 8-week-old *Gata4*^P-/-^ mice showed a normal histology except for a slightly weaker eosinophilic cytoplasm in acinar cells (**Fig. 1B**) and a non-statistically significant increase in lipomatosis (**Fig. 1C**). Expression of Ki-67, MCM4, cleaved-caspase 3, and γH2AX was similar in control and *Gata4*^P-/-^, indicating no major differences in proliferation, apoptosis and DNA damage (**Fig. 1D**). Similarly, acinar cell polarity was preserved, as MUC1 and E-cadherin were detected in mutually exclusive cell membrane domains (**Fig. 1E**). KRT19 was undetectable in normal acinar cells but was weakly detected in acinar cells of *Gata4*^P-/-^ mice (**Fig. 1D**). These results are unlike those found in 8-week-old *Gata6*^P-/-^ mice, where increased proliferation, apoptosis, and DNA damage, as well as altered cell polarity, were observed (Martinelli et al., 2013). In addition, *Gata4*^P-/-^ mice did not show spontaneous loss of the exocrine parenchyma during adulthood, in contrast with findings in *Gata6*^P-/-^ mice.

To identify molecular differences with higher resolution, we performed RNA-seq analysis of bulk pancreatic tissue from 8-10-week-old control and *Gata4*^P-/-^ mice and compared the results to the effects previously shown in *Gata6*^P-/-^ mice (Martinelli et al., 2013). Principal Component Analysis (PCA) showed that the samples from *Gata4*^P-/-^ mice cluster separately from both control and *Gata6*^P-/-^ samples, indicating that the global transcriptomic effects of embryonic GATA4 inactivation are distinct from those of GATA6 knockout (**Fig. 1F, Suppl. Fig. 1B, Suppl. Table 1**). These differences are further illustrated by the fact that the number of genes dysregulated upon *Gata4* inactivation is much lower than upon *Gata6* inactivation (**Fig. 1G, Suppl. Fig. 1C**) and few of them are concordantly dysregulated in *Gata4*^P-/-^ and *Gata6*^P-/-^ pancreata (**Suppl. Fig. 1D, Suppl. Table 1**). We then compared the regulatory pathways potentially affected by either *Gata4* or *Gata6* inactivation via Gene Set Enrichment Analysis (GSEA) with HALLMARK and REACTOME libraries and a signature of acinar genes (Masui et al., 2010). Unlike *Gata6* inactivation, associated with differential activity of a large number of pathways related to inflammation, cell cycle, ECM, and metabolism, *Gata4*^P-/-^ pancreata showed a much lower number of affected gene signatures (**Fig. 1H**, **Suppl. Fig. 1D, Suppl. Table 2)**. The gene set comprising digestive enzymes was the only one displaying similar changes in both *Gata4*^P-/-^ and *Gata6*^P-/-^ pancreata. In contrast, pathways related to collagen organization and TLR processing were dysregulated in opposite directions (**Suppl. Fig. 1D, Suppl. Table 2**).

**Supplementary Figure 1.**
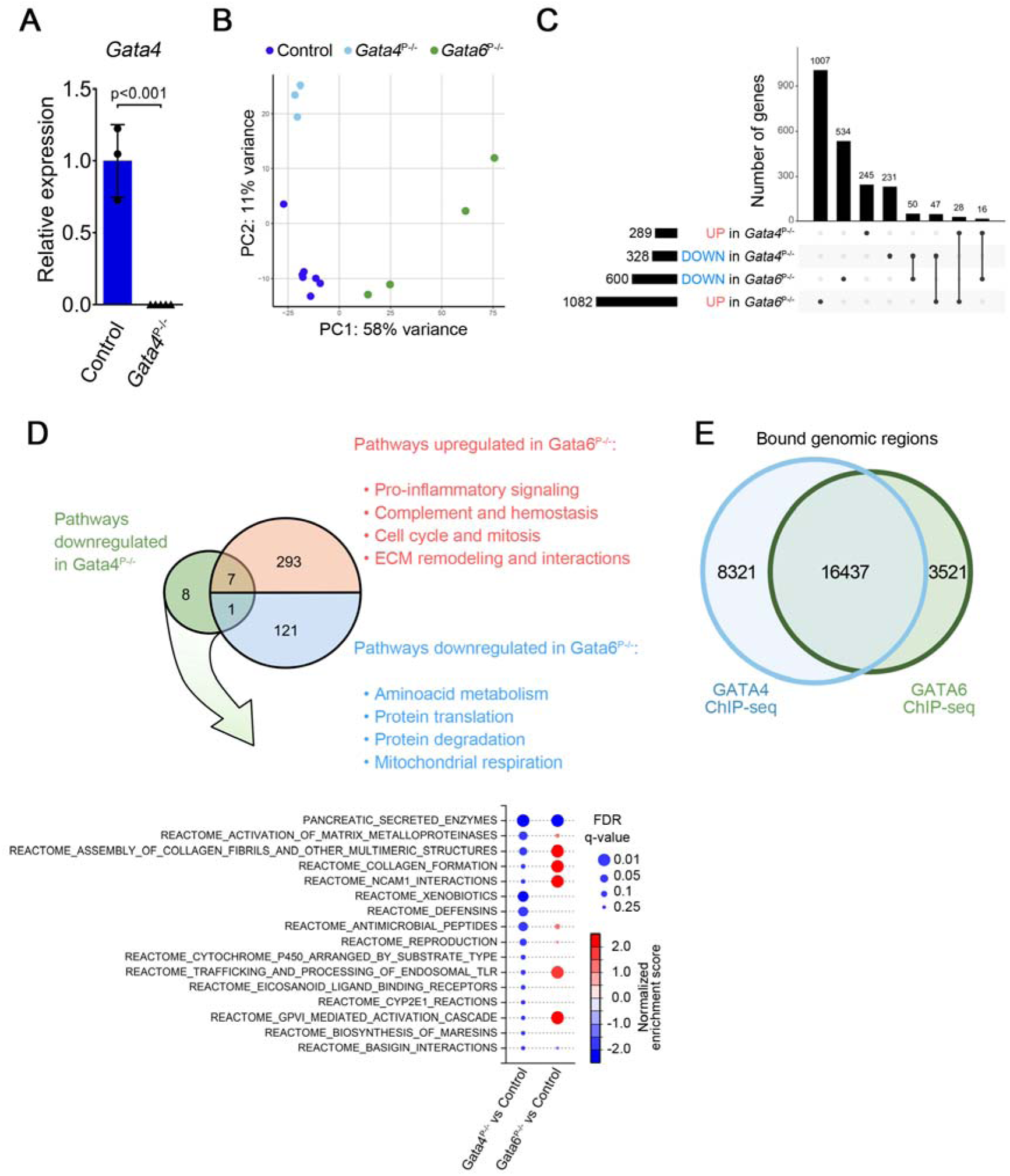
*Gata4* inactivation in the embryonic pancreas has a transcriptomic effect different from that of *Gata6* inactivation. **(A)** Normalized expression levels of *Gata4* mRNA in control (n=3) and *Gata4^P-/-^* mice (n=5) assessed by RT-qPCR**. (B)** PCA plot for RNA-seq data from control (n=7), *Gata4*^P-/-^ (n=3), and *Gata6*^P-/-^ (n=4) pancreata. **(C)** UpSet plot representation of the overlap between DEGs affected by inactivation of *Gata4* or *Gata6*. **(D)** Summary of GSEA results comparing transcriptomic signatures affected by *Gata4* and *Gata6* inactivation. Signatures with FDR q-value <0.25 are shown. **(E)** Venn diagram representing the number of genomic regions bound by either GATA4 or GATA6 in bulk pancreatic tissue using ChIP-Seq.

To identify genetic programs directly regulated by GATA4, we performed ChIP-Seq on chromatin from bulk normal pancreas tissue. Motif analysis confirmed ChIP specificity, with the canonical GATA consensus emerging as the most enriched motif. The top enriched motifs corresponded to established regulators of acinar differentiation, indicating that GATA4 and other pancreatic transcription factors co-occupy shared regulatory elements. (**Fig. 1I**). Approximately 34% of GATA4-bound regions localized near promoters (±1 or ±5 kb from the TSS) (**Fig. 1J**). We validated GATA4 binding at promoters of key acinar genes, including *Pnlip*, *Cpb1*, and *Try4*. Comparison with published GATA6 ChIP-seq data (Martinelli et al., 2013) showed that most binding sites are shared between the two factors, despite their distinct transcriptomic impact (**Suppl. Figure 1E, Suppl. Table 3**). We therefore conclude that the milder phenotypic changes observed in GATA4-deficient pancreas are secondary to its impact on downstream genes.

To determine whether *Gata4*^P-/-^ mice display a compromised response to damage, we induced a mild 7 hourly caerulein acute pancreatitis. *Gata4*^P-/-^ pancreata showed a slight increase in edema, acute inflammation, and vacuolization compared with control mice. (**Suppl. Fig. 2A**). *Gata4*^P-/-^ mice developed a lower rate of ADM but the differences were not statistically significant. *Gata4^-/-^* acinar cells can undergo metaplasia that is resolved after one week (**Suppl. Fig. 2A,B**). These findings are unlike those in *Gata6*^P-/-^ mice which had more damage and failed to recover properly from mild acute damage.

Altogether, we conclude that - unlike *Gata6* - *Gata4* is required for a normal pancreatic transcriptome but is largely dispensable for the maintenance of the exocrine pancreas, it partially overlaps with GATA6 function, and it is not absolutely required for the response to mild damage.

**Supplementary Figure 2.**
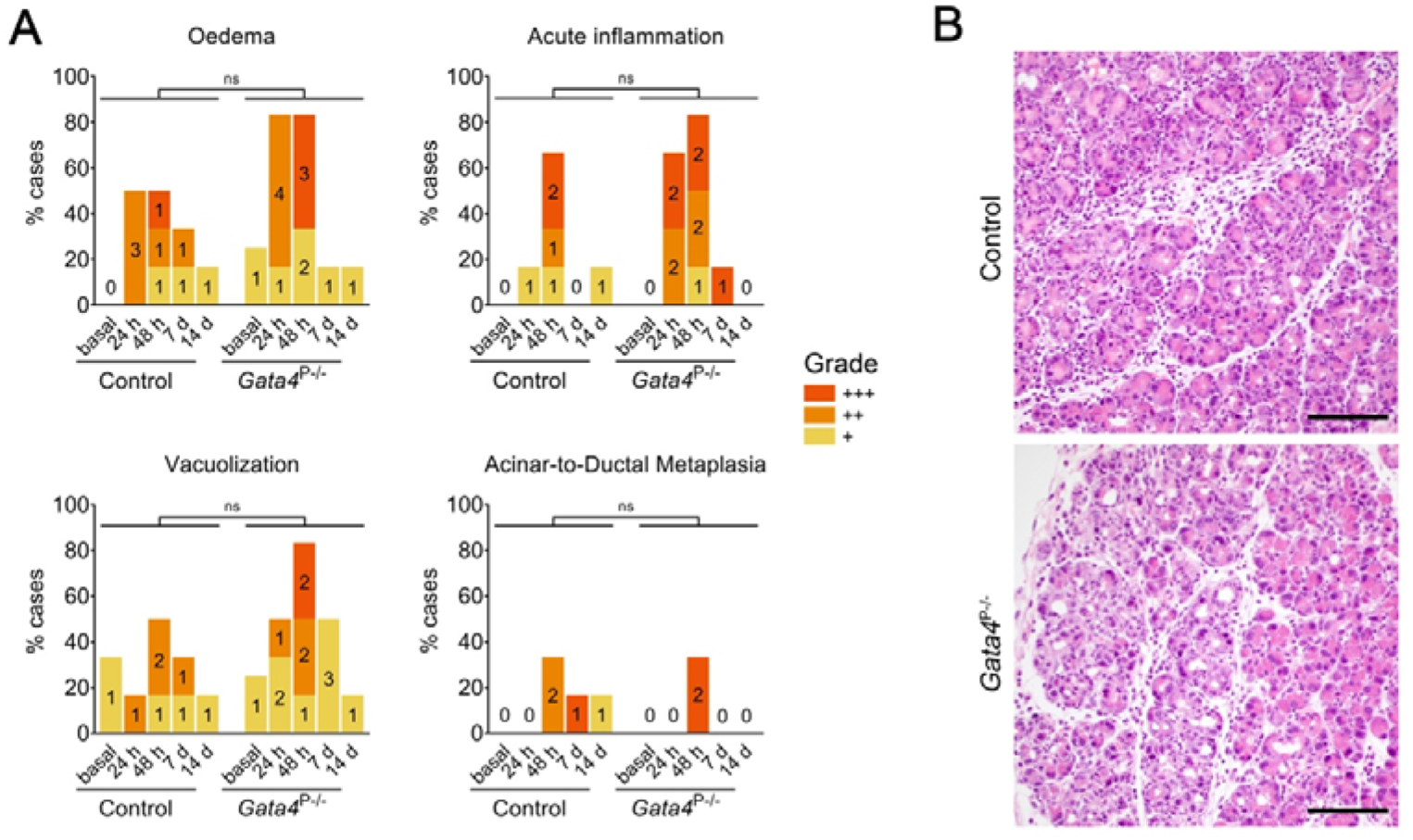
*Gata4^P-/-^* mice recover normally from acute pancreatitis. An acute pancreatitis was induced by 7 hourly caerulein injections. Pancreata (n=6/group) were harvested 24h and 48h post-caerulein administration (damage phase) and at days 7 and 14 (recovery phase). Control mice received saline and were included as basal state (n=3 for controls; n=4 for *Gata4*^P-/-^). **(A)** Damage was scored according to the presence of oedema, leukocyte infiltration, acinar vacuolization, and ADM, categorized as: 1=minimum; 2=mild; 3=moderate. **(B)** Representative H&E images (48 h) of mild and moderate ADM in *Gata4*^P-/-^ and control mice, respectively. Scale bar, 100 µm.

### *Gata4* deletion in *Kras*-mutant pancreata leads to PDAC in the absence of canonical precursor lesions

To assess the impact of *Gata4* deletion on mutant *Kras*-driven PDAC, we generated *Ptf1a*^+/Cre^ (control), *Ptf1a*^+/Cre^;*Kras*^+/LSL(G12Vgeo)^ (KC), and *Ptf1a*^+/Cre^;*Kras*^+/LSL(G12Vgeo)^;*Gata4*^lox/lox^ (KG4C) mice. Low-grade PanINs were almost absent in KG4C pancreata at all time points analyzed, in marked contrast with findings in KC mice (<40 weeks, P=0.003; 1 year, P<0.001; 1.5 years, P<0.001) (**Fig. 2A,B).** AFL were also markedly reduced in KG4C compared to KC mice at all timepoints [<40 weeks: 2/7 vs. 0/10 (P=0.154); 1 year: 1/23 vs. 10/14 (P<0.001); 1.5 years: 1/6 vs. 7/14 (P=0.325)] (**Fig. 2B**). This was accompanied by a significantly lower rate of tissue remodelling in KG4C pancreata. High-grade PanINs were infrequent in both KG4C and KC mice (**Fig. 2B**) and mucinous cystic lesions were not found in either strain. Despite the very low incidence of canonical preneoplastic lesions, the occurrence of PDAC was similar in KG4C vs. KC mice at all timepoints analyzed [<40 weeks: 1/10 vs. 2/17 (P=1.00); 1 year, 9/23 vs. 5/14 (P=1.00); 1.5 years, 3/6 vs. 7/14 mice (P=1.00)] (**Fig. 2A,B**). Extensive lipomatosis was observed in KG4C pancreata at <40 weeks (**Fig. 2C**), which became more prominent as mice aged, especially after 1 year.

**Figure 2.**
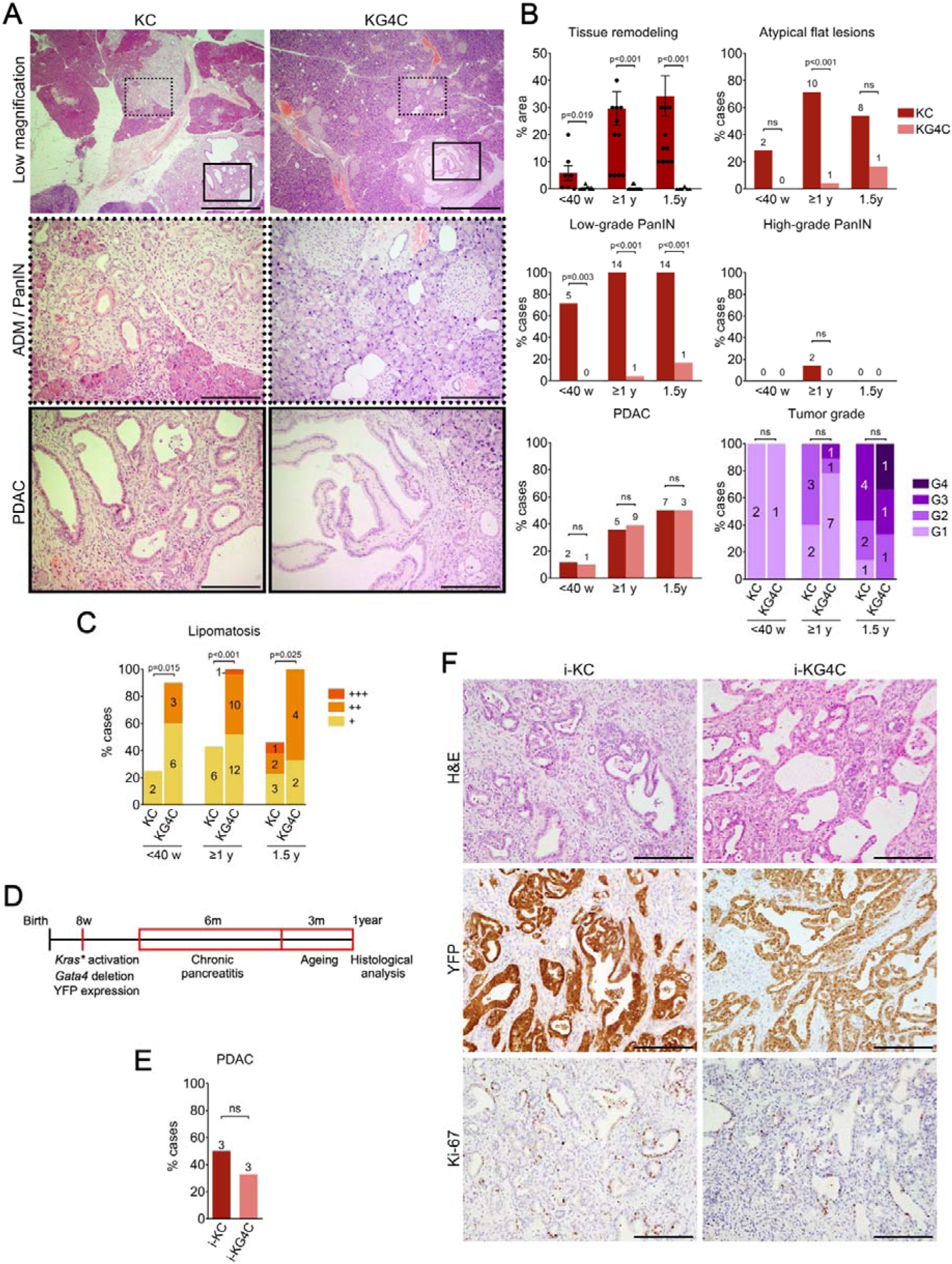
KG4C mice develop PDAC in the absence of PanINs. **(A)** Representative H&E images of the pancreas of 1-year-old KC and KG4C mice. Images of representative areas of ADM/PanIN (dotted line) and PDAC (solid line) at low (upper) and high magnification. Scale bars: 1 mm (low magnification) and 200 µm (high magnification). **(B)** Age-stratified quantification of tissue remodelling, low-grade PanIN, Atypical Flat Lesions (AFL), high-grade PanIN, and PDAC, including tumor grade. Tissue remodeling, low-grade PanINs, and AFL were prevalent In KC pancreata and almost absent in KG4C pancreata. In contrast, the incidence of PDAC was similar in the two groups. Number of KC mice analysed: <40w (n=10); 1y (n=14); 1.5y (n=14). Number of KG4C mice analysed: <40w (n=7); 1y (n=23); 1.5y (n=6). For tumor grade, only mice bearing tumors were considered. For KC mice at <40weeks, PDAC incidence was analysed in an extended series of 17 cases. **(C)** Increased lipomatosis in KG4C pancreata. **(D)** Experimental set-up for mutant *Kras* activation in adult mice (inducible model): two weeks after TMX administration, a chronic pancreatitis was induced by administering one supramaximal dose of caerulein daily for 6 months. Three months later, mice were sacrificed for histological analysis. **(E)** Quantification of tumor incidence in i-KC and i-KG4C mice showing no significant differences between the two groups (P=0.622). **(F)** Representative images of tumors from i-KC and i-KG4C mice. IHC detection of YFP and Ki-67 in tumors indicates an acinar origin and similar proliferation rates, respectively. P>0.05 (ns). Scale bar, 200 µm.

To assess the impact of acinar-specific *Gata4* deletion in the adult, we generated *Ptf1a*^+/CreERT2^; *Kras*^+/LSL(G12Vgeo)^;*Gata4*^lox/lox^; *Rosa26*^YFP/YFP^ and *Ptf1a*^+/CreERT2^; *Kras*^+/LSL(G12Vgeo)^ mice (termed i-KG4C and i-KC, respectively). Mice received tamoxifen (TMX) at 8 weeks, followed by one supramaximal dose of caerulein (0.25 mg/kg) daily for 6 months to induce chronic damage (**Fig. 2D,E**). The prevalence of tumors at 1 year in both mouse strains was not significantly different (3/9 - tumors in i-KG4C mice; 3/6 – tumors in i-KC mice; P=0.622) (**Fig. 2E**). Tumors from both i-KC and i-KG4C mice showed ductal morphology and widespread YFP expression, demonstrating their acinar origin (**Fig. 2F**). Again, low-grade PanINs were present in all i-KC (6/6) mice but not in i-KG4C pancreata (0/9) (**Fig. 2F** and data not shown).

These results indicate that upon *Gata4* deletion, either in all pancreatic epithelial cells or in adult acinar cells, mice carrying a mutant *Kras* allele develop PDAC in the absence of canonical precursor lesions.

### *Gata4* acts as a tumor suppressor and cooperates with *Trp53* to drive undifferentiated PDAC

To further address the effect of *Gata4* deletion on pancreatic carcinogenesis, we generated *Ptf1a*^+/Cre^;*Kras*^+/LSL(G12Vgeo)^;*Trp53*^lox/lox^ and *Ptf1a*^+/Cre^;*Kras*^+/LSL(G12Vgeo)^;*Trp53*^lox/lox^*;Gata4*^lox/lox^ (KPC and KPG4C, respectively). At humane endpoint, the incidence of PDAC was 100% in both mouse strains. The area affected by ADM, the number of AFL, and the occurrence of PDAC were also not significantly different. We found a non-statistically significant higher prevalence of cystic structures in KPC than in KPG4C pancreata (**Fig. 3A**). The survival of KG4C and KPG4C mice was significantly shorter than the corresponding *Gata4*-proficient controls (P=0.02 and P<0.001, respectively) (**Fig. 3B**). Similar results were obtained upon heterozygous *Trp53* deletion (P=0.002).

**Figure 3.**
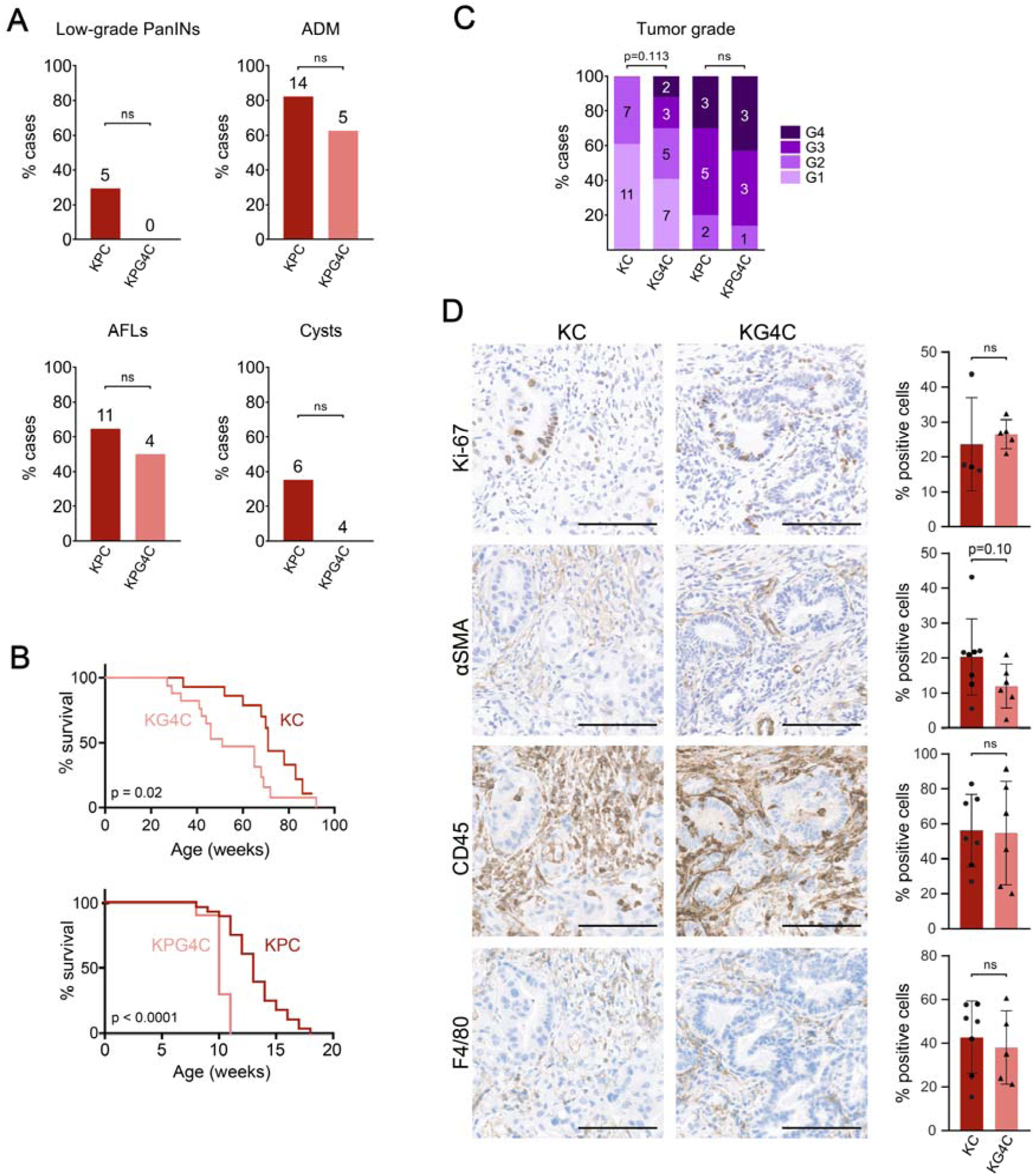
GATA4 acts a PDAC tumor suppressor in mice. **(A)** Prevalence of pre-neoplastic lesions and tumors in KPC and KPG4C pancreata at humane endpoint. Sample size: KPC (n=17), KPG4C (n=8). **(B)** Kaplan-Meier survival plots of KC (n=16) and KG4C (n=23) mice showing a significantly lower survival in the latter (Log-rank test, P=0.02) (upper panel). Kaplan-Meier survival plots of KPG4C (n=10) and KPC mice (n=28), showing a shorter survival in the absence of *Gata4* (P<0.001) (lower panel). **(C)** Distribution of tumors by grade. **(D)** Quantification and representative images of immunostainings for Ki-67, αSMA, CD45, and F4/80 in PDAC from KC and KG4C mice. Scale bar, 100 µm. Graphs show mean ± SD. Each dot represents an independent tumor. Two-sided Mann–Whitney test.

Lack of expression of GATA4 and/or GATA6 in patient tumors is associated with a basal phenotype and worse outcome (de Andres et al. 2023). KG4C tumors showed a tendency towards reduced differentiation, compared with KC tumors (P=0.113) (**Fig. 3C**), consistent with their more aggressive behavior (P=0.02). Similar results were obtained in KPG4C mice (**Fig. 3C**) and in murine PDAC arising in a variety of genetic backgrounds (**Suppl. Fig. 3**). As in human PDAC, GATA4 and GATA6 expression were highly correlated (r=0.63) and associated with grade (**Suppl. Fig. 3C,D**). Tumors from KG4C mice showed a modest increase in the percentage of Ki-67+ epithelial cells. There were no significant differences in the proportion of CD45+ tumor-infiltrating leukocytes or macrophages, but we found a non-significant tendency to reduced number of SMA+ cells in tumors from KPG4C mice (P=0.1), supporting a role for GATA4 in shaping the PDAC microenvironment (**Fig. 3D**).

Altogether, these results indicate that *Gata4* acts as a *bona fide* tumor suppressor and cooperates with *Trp53* to drive *Kras*-initiated PDAC progression.

**Supplementary Figure 3.**
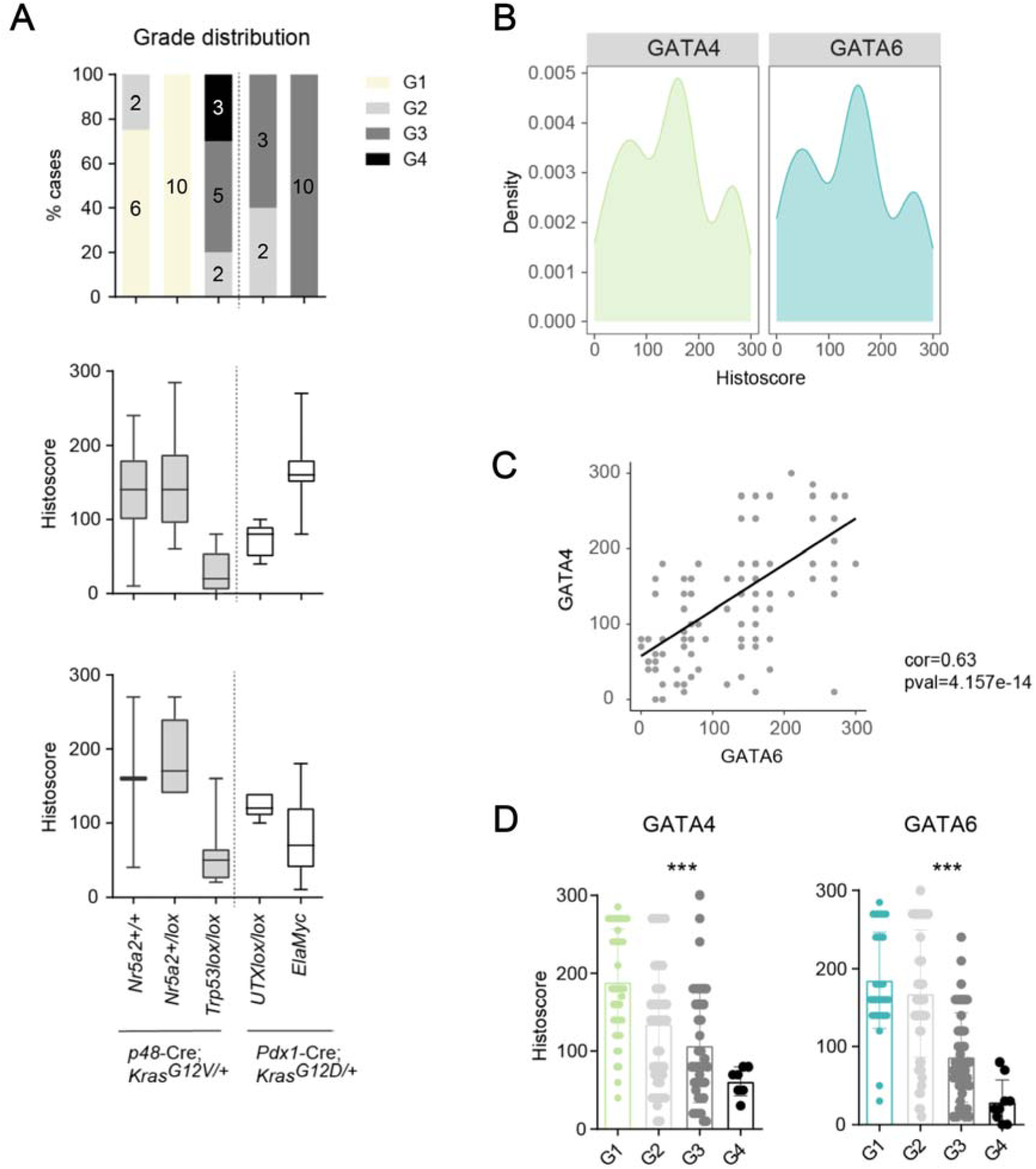
GATA6 and GATA4 expression correlate and are strongly associated with differentiation in mouse PDAC. **(A)** Top: Stacked bar plot showing tumor grade distribution. Number of mice per strain and group is displayed. Bottom: Box plots representing GATA4 and GATA6 histoscore in tumors from analyzed strains. Histoscore was calculated as intensity of staining (0-3) x % positive cells. **(B)** Density plot showing GATA histoscore distribution in all tumors analyzed (n=112). **(C)** Scatter plot showing a direct correlation between GATA4 and GATA6 histoscore across all samples (Pearson’s test). **(D)** Bar plots representing GATA4 and GATA6 histoscore classified by tumor grade. Graph shows mean ± SD. One-way ANOVA was used to calculate statistical significance. ***, P<0.001.

### PALes: putative mutant *Kras*-driven preneoplastic lesions arising in a *Gata4*-null context

The lower rate of ADM and lack of PanINs, AFLs, and IPMNs in KG4C and i-KG4C mice raises the possibility that an alternative route of PDAC development exists.

Histological analysis revealed the presence - at all time points analyzed - of unconventional metaplastic acinar lesions characterized by a pale cytoplasm with reduced eosinophilia and variable hyperplasia, morphologically different from ADM (**Fig. 4A-C**). We have designated them “PALes” for “<u>P</u>ale <u>A</u>cinar <u>Le</u>sions”. PALes were significantly more common in young (8-12-week-old) KG4C than in KC mice and their frequency increased transiently upon induction of an acute pancreatitis (**Fig. 4A,B**). PALes resemble ductular complexes, are surrounded by a normal acinar parenchyma, and are found both in KC and KG4C pancreata. **Table 1** summarizes the molecular features of PALes, compared to ADM and PanINs, in 8-12-week-old mice. In contrast to PanINs, PALes lack mucins (**Fig. 4C**). PALes showed low or undetectable expression of canonical acinar markers and robust expression of SOX9; expression of KRT19 was generally weaker in PALes than in ADM or PanINs (**Fig. 4D**). Ki-67 expression was reduced in PALes compared to PanINs (**Fig. 4E**); p21 and p16 were detected in 20% of PanINs and were broadly detectable in PALes. DCLK1, a marker of regenerative stem cells in vitro and in vivo, was expressed with a scattered pattern both in PALes and in PanINs from KC pancreata (**Fig. 4E**), consistent with previous reports (Westphalen et al., 2016). Cleaved caspase 3 expression was similar in PALes and PanINs. F4/80+ macrophages were significantly decreased in PALes, compared to PanINs. The trend was similar for neutrophils, while T or B cell infiltration was similar in both types of lesions (**Fig. 4F** and data not shown). PALes typically showed a reduced stromal reaction, compared to PanINs. These results indicate that PALes have a distinct phenotype and specific microenvironmental features.

**Figure 4.**
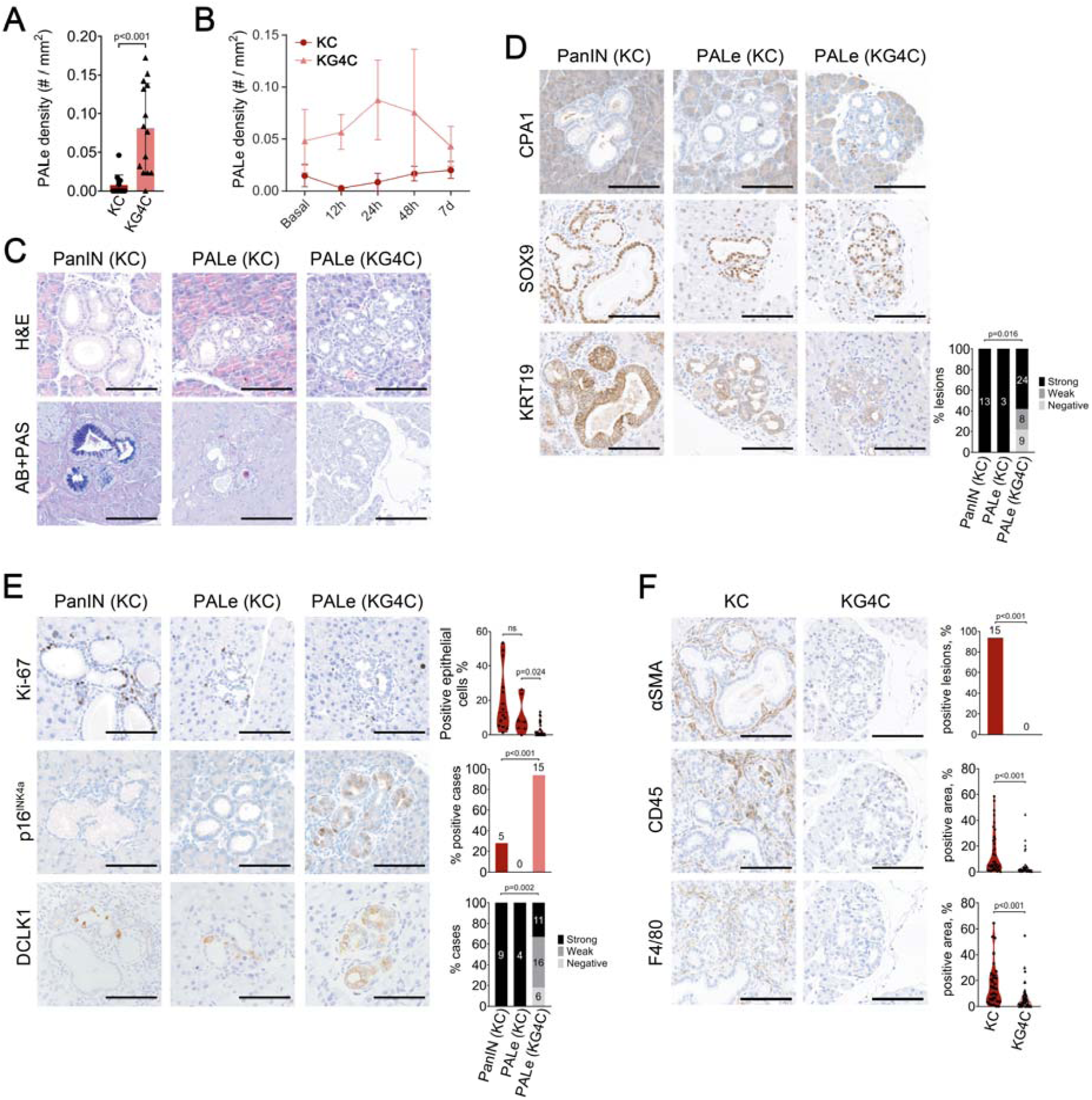
PALes are candidate mutant *Kras*-induced preneoplastic lesions in mice. **(A)** Prevalence of PALes in the pancreas of KC and KG4C mice. **(B)** Dynamics of PALe prevalence in KC and KG4C mice after a mild acute pancreatitis (n=4 for each genotype and time point). **(C)** Representative images of H&E and AB+PAS stainings (10 weeks). **(D)** Representative images of IHC analysis of expression of CPA1, SOX9, and KRT19 in PanINs and PALes (10 weeks) (n>4 mice/genotype). **(E)** Representative images and quantification of Ki-67, p16, and DCLK1 expression in epithelial cells from KC and KG4C mice (10 weeks). The number of lesions analyzed is shown in each graph (n=>3 mice/genotype). **(F)** Representative images and quantification of αSMA, CD45, F4/80, expression in lesions from 10-week-old KC and KG4C pancreata (n=6/genotype). Exact Fisher’s test and two-sided Mann–Whitney test were used to calculate statistical significance. was used to calculate statistical significance. Scale bar, 100 µm.

**Table 1.** Comparison of the molecular phenotype of ADM, PanINs, and PALes*.

| **Marker** | **ADM** | **PanIN** | **PALe (KC)** | **PALe (KG4C)** |
| --- | --- | --- | --- | --- |
| CPA | - | - | - | - |
| SOX9 | +++ | +++ | +++ | +++ |
| KRT19 | +++ | +++ | +++ | +++ |
| HNF1B | ++ | ++ | - | - |
| REG3B | ++ | +++ | - | - |
| Mucins | - | +++ | - | - |
| GKN | - | +++ | - | - |
| DCLK1 | ++ / + | ++ | ++ | +/- |
| Ki-67 | + | ++ | + | +/- |
| p16 | + /- | +/- | - | +++ |
| SMA | + | +++ | - | - |
| CD45 | + | +++ | + | + |
| F4/80 | ++ | +++ | + | + |

\* Summary information compiled from the literature and this manuscript

To assess whether *Gata4* inactivation had an impact on the composition of the microenvironment of PALes, we compared lesions from 8-12-week-old KC and KG4C pancreata. The proportion of epithelial Ki-67+ cells was significantly lower in KG4C PALes compared to KC PALes and PanINs (**Fig. 4E**). KG4C PALes showed a significantly lower number of αSMA+ cells, CD45+ cells, and F4/80+ cells than KC PanINs (**Fig. 3D**, **Fig 4F**), indicating a role for GATA4 in shaping the epithelial microenvironment.

### GATA4 is required for an early, mutant KRAS-driven, transcriptional inflammatory program

To acquire insight into the mechanisms operating upon *Gata4* deletion, we performed bulk RNA-seq of 8-10-week-old control (*Ptf1a*^+/Cre^), KC, and KG4C pancreata. At this age, the pancreas is histologically normal. PCA showed consistent transcriptomic differences between all three experimental groups (**Fig. 5A**). Seven clusters of differentially expressed genes (DEGs) were identified (**Fig. 5B, Suppl. Tables 4,5**). Clusters 1 and 2 included genes up-regulated in mutant *Kras* pancreata, the expression of which was reduced upon *Gata4* deletion. These gene sets were enriched in immune cell markers (e.g., *Cd4, Cd8, Cd19*), chemokines and their receptors (e.g., *Ccl5, Cxcl9, Cxcr5*), and *Reg* family members (*Reg1, Reg3a, Reg3b*). Cluster 3 included transcripts that were minimally affected by mutant *Kras* alone but the expression of which was markedly up-regulated when *Gata4* was concomitantly deleted; this gene set was enriched in ductal cell markers (e.g., *Onecut1, Sox9*). Other clusters included genes down- (e.g., cluster 4, *Amy2, Prss3*) or up-regulated (e.g., cluster 5, *Col2a1*) upon mutant *Kras* expression, as well as two clusters containing a small number of genes (6 and 7).

**Figure 5.**
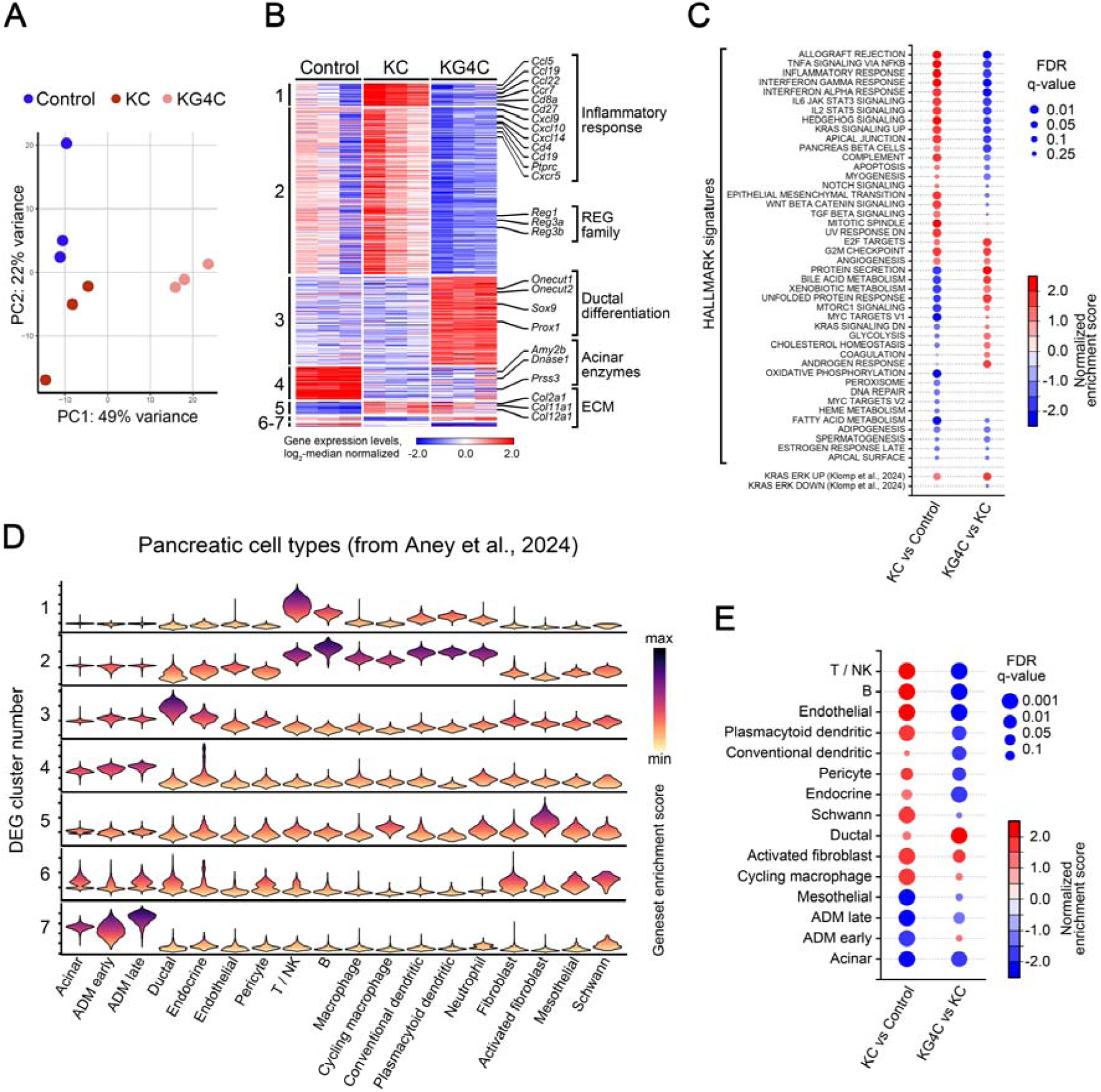
GATA4 is required for the activation of the transcriptional inflammatory programs induced by mutant *Kras*. (A) PCA plot of RNA-seq data from C (n=3), KC (n=3), and KG4C (n=3) pancreata. (B) Heatmap representation of DEGs across the three experimental groups. Distinct clusters of DEGs are indicated; individual genes and processes in which they participate are highlighted. (C) GSEA of differences in the enrichment of MSigDB HALLMARK and KRAS-ERK signatures (Klomp et al., 2024) in two-way comparisons. (D) Violin plot representation of enrichment scores for each cluster of DEGs across pancreatic cell populations identified in Aney et al., 2024. (E) Changes in GSEA-estimated abundance of cell types in bulk pancreas. Cell type signatures derived from Aney et al., 2024; cell types showing no significant differences are omitted.

Mutant *Kras* activation was associated with a significant up-regulation of pathways related to KRAS signaling (e.g., *Etv5*, *Dusp6*, *Ephb2*), cell cycle (e.g., *Aurka*, *Ube2c*, *Cenpf*), and inflammation (multiple genes from the *Ccl*, *Cxcl*, and *Il* families) (**Fig. 5C**, **Suppl. Table 6**). The inflammation-associated signatures, but not the cell cycle pathways, were suppressed upon *Gata4* deletion. Surprisingly, the activity of refined KRAS-ERK pathway signatures (Klomp et al., 2024) was further enhanced in KG4C pancreata, suggesting that canonical KRAS-ERK signaling alone is not sufficient to drive the pro-inflammatory changes (**Fig. 5C**). We next explored the associations between identified DEGs with epithelial and non-epithelial cell types present in the murine pancreas, as defined using a recent scRNA-seq dataset (Aney et al., 2024) (**Suppl. Fig. 4A, Suppl. Table 7**). The enrichment scores for the genes included in the clusters defined above indicated that Clusters 1 and 2 are largely attributable to myeloid, T-, NK-, and B cells. In contrast, DEGs in clusters 3 and 4/7 are mainly attributable to the ductal and acinar cell populations, respectively. Genes from cluster 5 were expressed in multiple cell populations but were specifically enriched in activated fibroblasts (**Fig. 5D**, **Suppl. Fig. 4B**). GSEA with cell type-specific signatures (**Fig. 5E**, **Suppl. Tables 7,8**) further supports the notion of a GATA4-dependent T-, NK-, and B-cell infiltration. On the other hand, the mutant *Kras*-driven increase in activated fibroblasts and cycling macrophages was not reverted upon *Gata4* inactivation (**Fig. 5E**). These findings were verified using canonical immune cell markers and the bulk RNA-seq data (**Suppl. Fig. 4C**).

To pinpoint the contribution of pancreatic epithelial cells to these expression changes we performed RNA-seq of sorted epithelial cells from 8-12-week-old control (n=4), KC (n=3), and KG4C (n=5) mice (**Suppl. Table 9**). Genes included in clusters 1, 2, 5, and 6 were not differentially expressed in epithelial cell populations, further supporting that the DEG patterns observed in bulk pancreatic tissue captured non-cell-autonomous changes within the immune (clusters 1,2) and fibroblast (clusters 5,6) compartments. On the other hand, cluster 3 genes followed a similar pattern of changes in bulk tissue and sorted epithelial cells, supporting a role of GATA4 in the maintenance of acinar over ductal lineage identities (**Suppl. Fig. 4D,E, Suppl. Tables 9-11**). Altogether, these findings indicate that GATA4 is selectively required for an early, mutant *Kras*-driven tissue remodeling state and inflammatory cell infiltration in the pancreas.

**Supplementary Figure 4.**
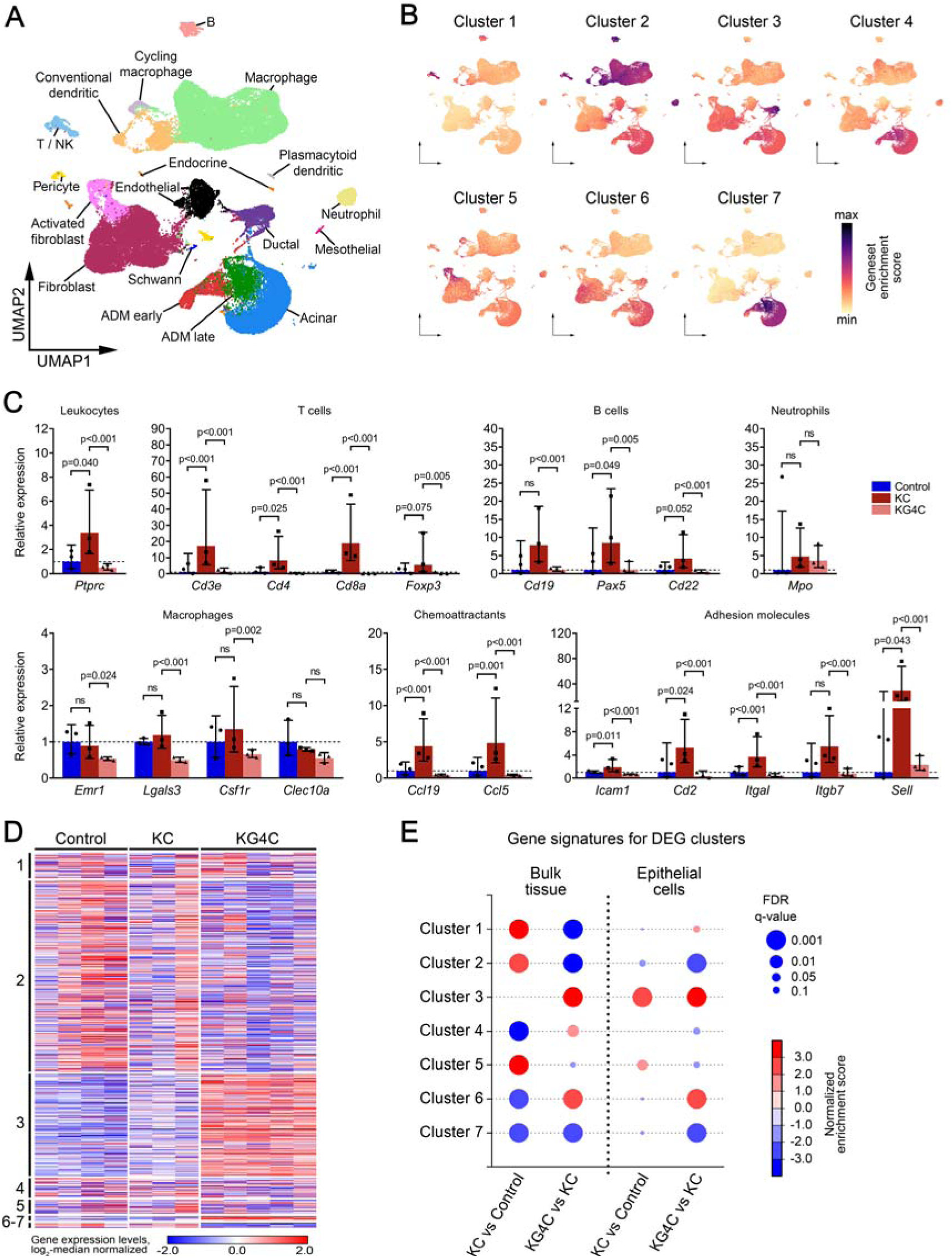
Inflammatory programs up-regulated in mutant *Kras-*expressing pancreata are largely attributable to immune cells. (A) UMAP representation of cell populations in normal pancreas (Aney *et al*., 2024). (B) Enrichment for each cluster of DEGs projected across pancreatic cell populations. (C) Relative expression levels of canonical immune cell markers in bulk pancreas across genotypes. (D) Heatmap representation of expression levels of DEGs corresponding to the clusters shown in Figure 5B in isolated pancreatic epithelial cells. (E) GSEA of differences in cluster 1-7 gene signature enrichment in bulk pancreas tissue and isolated epithelial cells.

### IL-17 rescues ADM - but not PanINs - in KG4C mice

ADM/ductal complexes have been shown to precede PanINs in mouse pancreatic carcinogenesis. We hypothesized that GATA4 controls the expression of a set of cytokines that are up-regulated in epithelial cells upon mutant *Kras* activation to orchestrate an inflammatory response and acinar reprogramming (ADM, ductal complexes) response, leading to PanINs. As shown above, macrophage density - unlike neutrophil, T or B cell density - was lower in PALes from KG4C mice (**Fig. 4F** and not shown), suggesting a defect in macrophage recruitment in the absence of *Gata4*.

To identify candidates, we queried RNA-seq data from KC and KGC4 pancreata, primary isolated acinar cells, and FACS-isolated epithelial cells, as well as pancreatic GATA4 ChIP-Seq data (**Suppl. Tables 3,4,9**). We focused on the KEGG Cytokine-cytokine receptor interaction geneset (KEGG mmu04060) which is strongly upregulated in KC pancreata and suppressed upon *Gata4* inactivation (**Suppl. Fig. 5A**). Leading edge analysis identified 15 genes that were statistically significantly overexpressed in “KC vs Control” and suppressed “KG4C vs KC” comparisons (**Suppl. Fig. 5B**). *Cxcl12* and *Cxcr4* were also identified as direct GATA4 targets in ChIP-seq and ChIP-qPCR (**Suppl. Fig. 5C,D, Suppl. Table 3**). In addition, LC-MS analysis of the secretome of primary acinar cultures identified REG3B as the most downregulated secreted protein (out of 34 down-regulated proteins) in KG4C acini (FDR q-value<0.1) (**Suppl. Fig. 5E, Suppl. Table 12**), recapitulating the RNA-seq findings.

Both *Cxcl12* and *Reg3b* were expressed at moderate-high levels in bulk pancreas and isolated primary acini/epithelial cells of KC mice, were down-regulated upon *Gata4* inactivation in KG4C mice and freshly isolated acini, and their regulatory regions were bound by GATA4 (**Suppl. Fig. 5C-E,G**). CXCL12 regulates macrophage recruitment via its receptor, CXCR4 (Kim et al., 2014; X. Li et al., 2019). To examine whether CXCL12 promotes PanIN formation, we induced a mild acute pancreatitis to 15-week old KC mice to amplify PanIN development and administered the CXCR4 antagonist Plerixafor (5mg/kg i.p., three times a week for 10 weeks) (Aboumrad et al., 2007). The biological activity of Plerixafor was demonstrated by the higher WBC count in treated mice (p<0.01). Nevertheless, PanIN quantification at endpoint did not reveal significant differences in PanIN formation between Plerixafor-treated vs control mice (p=0.923).

REG3B is up-regulated during acute pancreatitis-associated ADM (Keim et al., 1984) and is involved in tissue macrophage recruitment (Lörchner et al., 2015) and polarization (Gironella et al., 2013). It was strongly expressed in KC mice surrounding areas of metaplasia and was absent from KG4C pancreata (**Suppl. Fig. 5H**). Interestingly, *Reg3b* knockout mice do not develop PanIN in a mutant *Kras* context (Loncle 2015). GATA4 regulates *Reg3b* in intestinal cells (Lepage et al., 2015) and our ChIP-Seq data identified a GATA4 binding site in an enhancer controlling the expression of *Reg* genes (Cuadrado et al., 2015) that was confirmed by ChIP-qPCR (**Suppl. Fig 5G**), suggesting direct regulation. *Ccl5* and *Ccl19* were two additional candidates identified in the KEGG Cytokine-cytokine receptor interaction geneset (KEGG mmu04060). However, they are expressed at very low levels in pancreatic tissue and isolated epithelial cells (**Suppl. Tables 4,9**), are not bound by GATA4, and expression did not differ between KG4C and KC acini. Therefore, we conclude that their expression changes in bulk pancreatic tissue likely arise from epithelial cell-driven effects on the immune cell compartment.

**Supplementary Figure 5.**
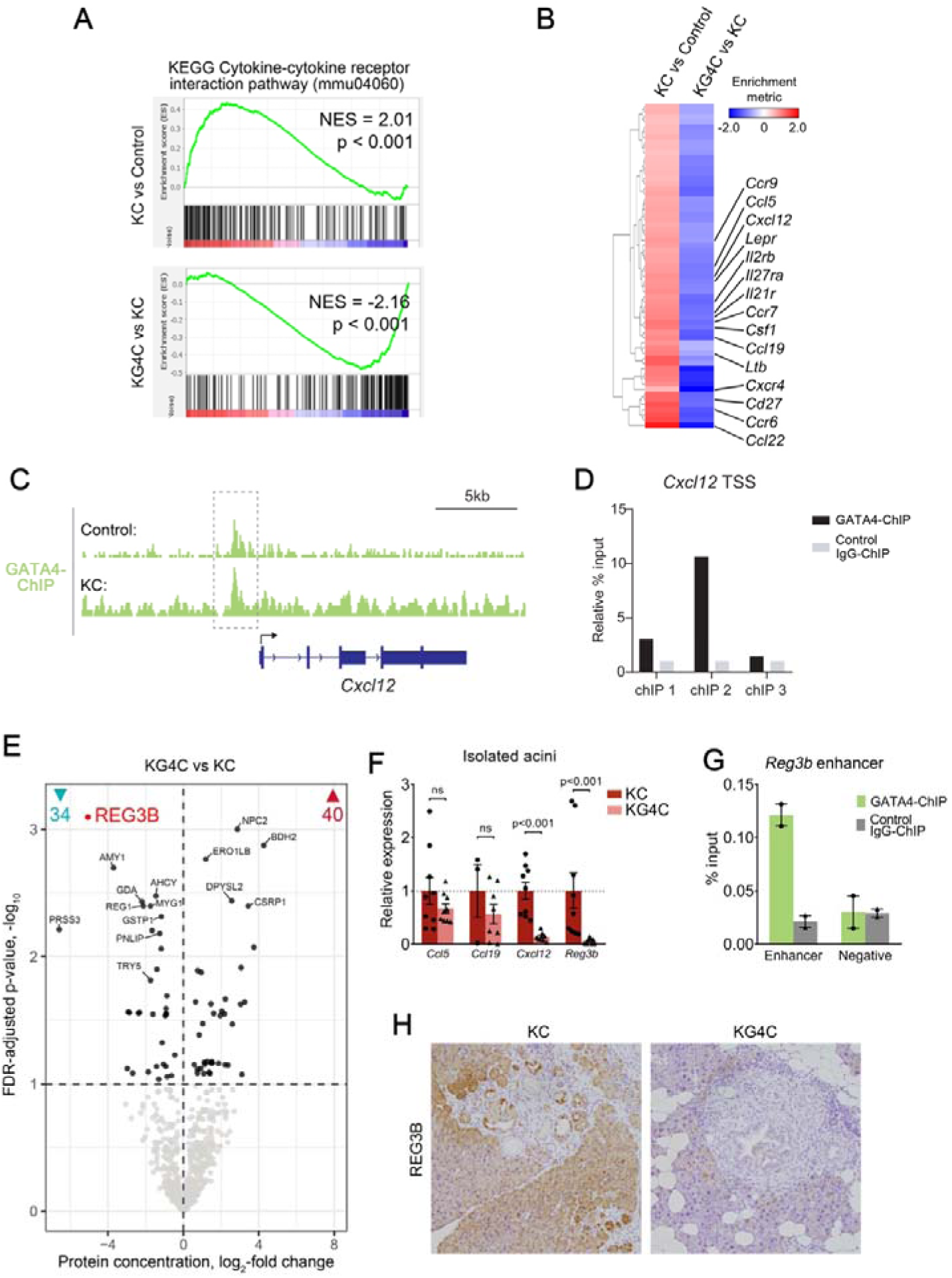
Candidate GATA4-dependent regulators of premalignant lesions and PDAC in *Kras*-mutant pancreas. (A) GSEA for KEGG Cytokine-cytokine receptor interaction geneset (mmu04060) in bulk pancreas tissue. (B) Leading edge analysis for genes from the mmu04060 geneset. Statistically significant DEGs are highlighted. (C-D) GATA4 binding to the *Cxcl12* promoter as shown by UCSC tracks (C) and validated by ChIP-qPCR (D). (E) Volcano plot representation of differential protein abundance in the secretome of KC and KG4C primary acinar cells. (F) Expression levels of selected cytokines in isolated acini from KC and KG4C mice. Two-tailed Student’s t-test performed on log-transformed values. (G) ChIP-qPCR validation of GATA4 binding to the enhancer region of *Reg3b*. (H) Immunostaining for REG3B in pancreatic tissue of KC and KG4C mice.

The histological and transcriptomic analyses suggested that GATA4 loss is associated with altered acinar-ductal transitions. To assess this defect, we turned to a functional ADM assay using Matrigel cultures (Miyamoto et al., 2003): *Gata4*^-/-^ acinar cells showed a dramatically reduced capacity to undergo ADM in vitro (P<0.001) (**Fig. 6A**). Several cytokines, including IL-17, IL-6, IL-1b and EGF, have been shown to promote ADM (Ardito et al., 2012; McAllister et al., 2014). As expected, recombinant EGF, IL-17, and IL-6 increased the ADM response in control cells; IL17 almost completely rescued the defect observed in *Gata4*^-/-^ acinar cells. Other cytokines (IL-1b, IL-2, IL-4, IL-10, CCL5, CXCL12, and REG3B) did not have a significant effect on ADM/cyst formation in control or *Gata4*^-/-^ cells (**Fig. 6A**). Acini from mice carrying a mutant *Kras* allele spontaneously formed a ca. 4-fold higher number of ADM/cysts compared to wild type acini (**Fig. 6B**). KG4C cells showed very reduced ability to form ADM/cysts in basal conditions (P=0.008) and IL-17 almost completely rescued this defect; EGF had a more modest effect (**Fig. 6B**). These results indicate that, regardless of *Kras* status, *Gata4* inactivation results in reduced *in vitro* ADM and that IL-17 rescues this defect.

**Figure 6.**
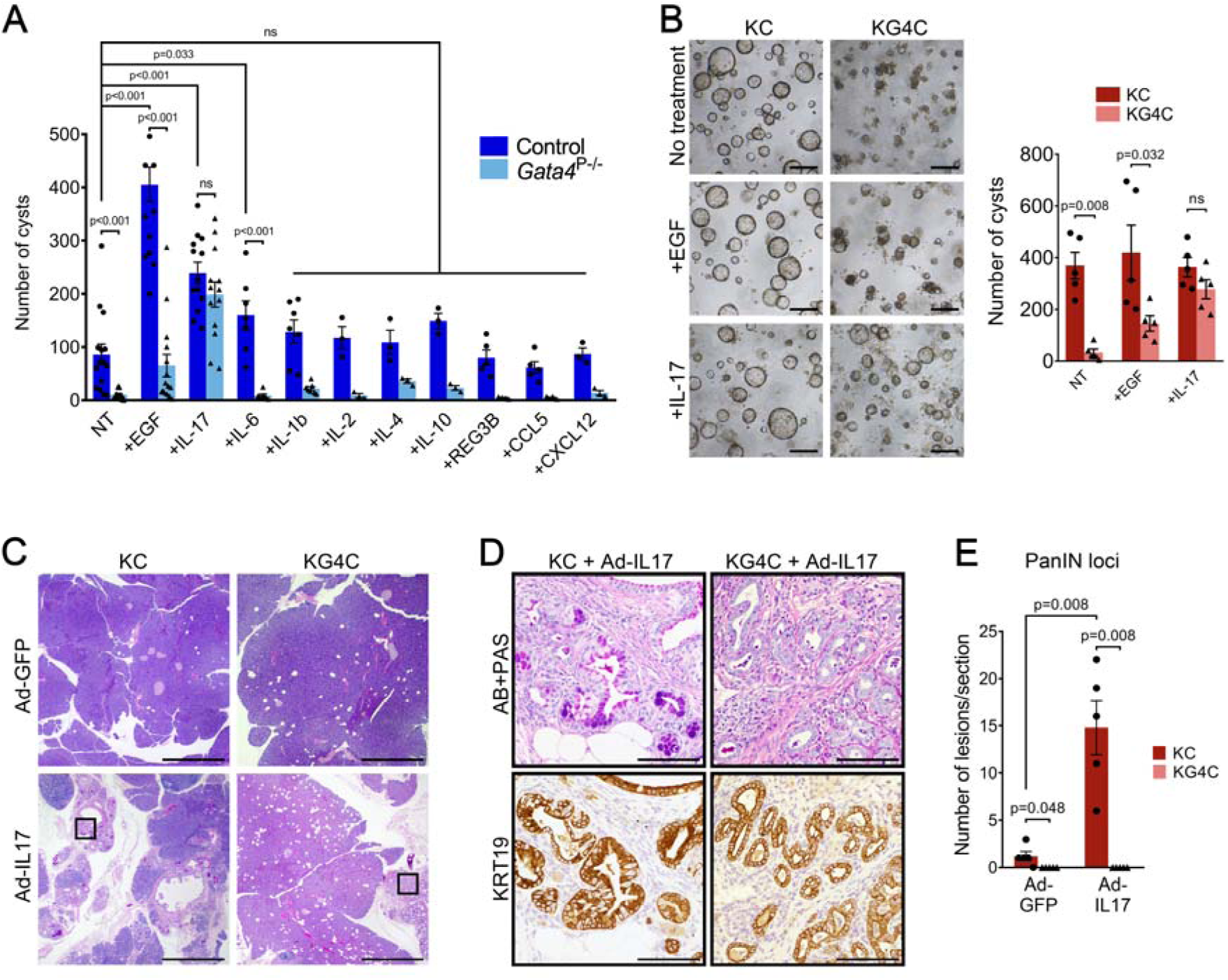
GATA4 is required for efficient ADM *in vivo* and *in vitro*. **(A)** Acinar cells from control and *Gata4*^P-/-^ mice were embedded in Matrigel and assayed for cyst formation, a surrogate of ADM: in the absence of GATA4, there is reduced ADM. This phenotype is rescued by IL-17 and, to a lesser extent, by EGF. Experiments/condition: control, EGF, IL-17 (n=13); IL-6, IL-1b (n=7); IL-2, IL-4, IL-10, REG3B, CCL5, CXCL12 (n=3-5). Two-sided Mann-Whitney test. **(B)** Phase microscopy images showing the effect of EGF and IL.17 on formation of cystic structures (n=5/group) (left panel). Scale bar, 200 µm. Cyst quantification (right panel). Two-sided Mann-Whitney test. **(C-E)** KC and KG4C mice were orthotopically injected with adenoviruses expressing either GFP or IL17 (10^9^ PFU/mice) (n=5/group). Mice were sacrificed 6 weeks later and the presence of low-grade PanINs was assessed using PAS/AB staining. **(C)** Representative H-E images. Scale bar, 1 mm. High magnification of insets shown in panel (**D**). **(D)** Low-grade PanINs were absent from KG4C pancreata both in control (Ad-GFP) and Ad-IL17-transduced mice. Scale bar, 100 µm. **(E)** Quantification of low-grade PanINs, showing a significant increase in KC mice injected with Ad-IL17 compared to Ad-GFP. Two-sided Mann-Whitney test.

Ectopic expression of IL-17 driven from adenoviruses (Ad) injected orthotopically in the pancreas of mice carrying a mutant *Kras* allele induces PanINs (McAllister et al., 2014). To determine whether IL-17 could rescue ADM and PanIN formation in the absence of GATA4, we injected Ad expressing either IL17 or GFP into the pancreas of 10-week-old KC and KG4C mice. Six weeks later, histological analysis showed that the number of PanINs was significantly higher in KC mice receiving Ad-IL17, compared to Ad-GFP (P=0.008) (**Fig. 6C-E**). In stark contrast, Ad-IL17 administration to KG4C mice resulted in increased metaplastic KRT19+ non-mucinous lesions, but PanINs remained undetectable (compared to KC mice, P=0.048 for Ad-GFP, and P<0.008 for Ad-IL17 inoculation) (**Fig. 6C-E**). These findings indicate that, in the absence of GATA4, ADM and PanIN formation is compromised but only the former is rescued by IL-17.

Collectively, these findings show that GATA4 directly controls the activation of pro-inflammatory factors in *Kras*-mutant acinar cells, which functionally contribute to canonical ADM formation and - more broadly - uncover a distinctive GATA4-dependent program that functionally uncouples PanIN and PDAC development.

## DISCUSSION

Understanding the early steps involved in PDAC development is critical to reduce its incidence and mortality. GEMMs have provided invaluable insight in these processes, which can hardly be analyzed in humans: for example, GATA proteins have shown the critical role of GATA6 in the differentiation of acinar cells and in the maintenance of the classical PDAC program. While both GATA6 and GATA4 are expressed at high levels in acinar cells, only the role of GATA6 has been explored using GEMMs. Here, we show that both genes play distinct roles in acinar homeostasis in the normal pancreas and in pancreas cancer initiation. We provide provocative evidence indicating that - in mice - PDAC can develop in the absence of all canonical preneoplastic lesions so far identified in humans or mice. Yet, they share roles in the maintenance of PDAC cell identity. Furthermore, we show that a GATA4-dependent, mutant KRAS-driven inflammatory program plays a more critical role in the development of PanINs than in the direct promotion of PDAC.

GATA4 and GATA6 have fundamentally different roles in the mouse pancreatic homeostasis and carcinogenesis. GATA6 is more critical than GATA4 in the maintenance of acinar and pancreatic tumor cell identity (Martinelli et al., 2013) (**Suppl. Table 3**). Both in *Kras*-wild type and -mutant contexts, *Gata4* and *Gata6* inactivation in the pancreas had opposite effects on the activity of inflammatory pathways and PanIN development: GATA4 is required for the pro-inflammatory effects of mutant KRAS whereas GATA6 has a suppressive role. These differential effects might be mediated by AP1 proteins that have been shown to participate in a pre-inflammatory pancreatic phenotype and in pancreatic carcinogenesis (Cobo et al., 2018; Li et al., 2024). Unlike *Gata4*, *Gata6* inactivation in *Kras*-mutant pancreata was associated with a significant up-regulation of inflammatory pathways (**Suppl. Fig. 1**) and an increase in PanINs, strongly supporting the role of inflammation in PanIN formation. Importantly, our work in mice shows that GATA4 and GATA6 retain, and share, a role in the maintenance of classical identity of PDAC as already highlighted by our prior work using patient samples (de Andrés et al., 2023).

The currently accepted model of PDAC development, derived from studies using GEMMs, posits that ADM is a precursor of low-grade PanINs which in turn are the major precursors of PDAC initiated in embryonic pancreatic epithelium or in adult acinar cells. A major tenet in pancreatic carcinogenesis is the role of inflammation as a promoter of PanINs and PDAC (Flandez et al., 2014; Guerra et al., 2007). Single-cell genomic analyses have shown that mutant Kras and inflammation cooperate to generate unique cancer-prone epigenomic states that foster tumor development (Alonso-Curbelo et al., 2021). We show that the mutant KRAS-dependent and GATA4-mediated pro-inflammatory changes reflect paracrine effects on haematopoietic cells. Beyond its impact on tissue remodelling, this phenotype was associated with lack of PanIN development but was permissive to the events leading to PDAC. Several studies have used low-grade PanIN development as a surrogate of PDAC but did not extend the work to analyze malignant tumor occurrence (Hamidi et al., 2012; Kopp et al., 2012; A. L. Li et al., 2024; Loncle et al., 2015). Our findings raise a note or caution in the interpretation of this work, as we clearly show that PDAC can develop in the absence of such lesions.

We show that ADM an PanIN formation are mechanistically dissociated as IL-17 was able to rescue the former but not the latter, indicating that distinct processes underlie the generation of both types of premalignant lesions. Intringuingly, while IL-17 regulates the expression of REG3B, shown to be required for PanIN formation (Loncle et al., 2015), REG3B administration to KG4C mice was not sufficient to rescue PanINs (not shown), indicating that GATA4 controls a broader functional program.

Our finding that - in mice - PDAC can efficiently develop in the absence of PanINs led us to search for putative neglected precursors. We have identified PALes as candidate precursors in the pancreas of mice harboring mutant *Kras* activated either in the embryonic or in the adult pancreas. While we do not have conclusive evidence that they are PDAC precursors, these lesions have not been observed in our laboratory in >1000 pancreata lacking a mutant *Kras* allele (not shown). Importantly, a search for PALes in human pancreata by two teams of highly esteemed pancreatic cancer pathologists (I.E. and Laura Wood, Johns Hopkins University) did not identify their presence in human pancreata using standard H&E staining. This is somewhat similar to observations of ADM, which is prominent in mice harboring mutant *Kras* in the pancreas but not in humans, even in subjects at high risk of PDAC. These findings should be considered in the context of a number of pitfalls related to the current canonical model of PDAC development in humans:

i) low-grade PanINs are pervasive in the adult human pancreas, even in young subjects. It has been estimated that the adult pancreas can contain up to ca. 1000 PanINs and that the risk of progression to PDAC is in the range of 10^-6^ (Braxton et al., 2024);

ii) work from the Hopkins group shows that essentially all low-grade PanINs found in non-neoplastic pancreas contain *KRAS* mutations, as PDAC. However, there is no direct evidence - neither in mice nor in humans - that PDAC arises from PanINs. It is conceivable that mutant KRAS could lead - independently - to PanINs or to PDAC depending on the cell genomic/epigenomic context in which the mutations occur. This hypothesis acquires more plausibility considering that work in the last decade has shown that the human body is a mosaic of cells with mutations in a large number of genes that have a relatively neutral impact on cell phenotypes (Coorens et al., 2025). Furthermore, a distinct epigenetic program has been identified in early *Kras*-mutant epithelial cells that is mutually exclusive in PDAC vs. PanINs (Burdziak et al. 2023);

iii) high-grade PanINs could arise independently from low-grade PanINs and be true precursors of PDAC, though a careful analysis of high-grade PanINs is required considering the fact that most current knowledge emanates from the study of samples acquired from patients with PDAC (Hosoda et al., 2017). The Hopkins group has provided evidence that high-grade PanINs from subjects without PDAC do not frequently harbor genetic alterations that are pervasive in those from patients with PDAC, strongly suggesting that much of the knowledge acquired using these samples may correspond to intraductal tumor spread (Hosoda et al., 2017);

iv) recent work shows that, in mice, a discrete population of *Kras*-mutant progenitor-like cells that undergo p53 loss are likely precursors of PDAC (Reyes et al., 2026).

The functional dissection of *Gata4* inactivation in mouse carcinogenesis points to a critical revision of the current conceptualization of PDAC development and highlights the relevance of species-specific cross-analyses in understanding the mechanistic underpinnings of these early processes.

## Study limitations

- There is substantial evidence on the conservation of GATA protein functions in the pancreas in mice and humans; nevertheless, species-specific differences may exist.

- The timing of occurrence of *KRAS* mutations in human pancreas has not been formally demonstrated but it is generally thought that they appear in the adult pancreas. Most of the work reported here was performed using *Ptf1a-Cre* as a driver of mutant *Kras* activation in pancreatic epithelial cells. However, the main findings of our work were replicated using *Ptf1a-CreERT2* to activate mutant *Kras* in adult acinar cells.

- The precise mechanisms through which GATA4 impacts on inflammation and reprogramming are not identified, nor are the phylogenetic relationships between ADM and PanINs.

- The significance of PALes as true preneoplastic lesions, in mice or humans, remains to be established.

- A possible caveat in the interpretation of the data is that GATA4, being a transcriptional regulator, could directly influence mucin gene expression.

## Supporting information

Description in the main text file

## Acknowledgements

We thank the members of the Epithelial Carcinogenesis Group for discussion and criticism, K. Aney, R. J. MacDonald, S. Nissim, F. Notta, C. Salomó, and L. Wood for valuable contributions, and the Animal Facility, Histopathology, and Genomics Units of CNIO for technical support.

## Funding

This work was supported, in part, by grants from the Ministerio de Ciencia, Innovación y Universidades (MCIU), Madrid, Spain to FXR (grants SAF2015-70553-R, RTI2018-101071-B-I00, and PID2021-128125OB-I00). Work in the lab of SWL was supported, in part, by grant CA283378 from the National Cancer Institute. SWL is an HHMI investigator and the Geoffrey Beene Chair for Cancer Biology. FM was supported by a Ph.D.Fellowship from La Caixa Health Research Foundation. MPA was supported by a Ph.D. Fellowship from MCIU and by an EMBO Scientific Exchange Grant (number 8446). MC was supported, in part, by a Postdoctoral Fellowship from “Amigos del CNIO”. CNIO is supported by MCIU as a Centro de Excelencia Severo Ochoa (grant SEV-2015-0510).

## Data access

RNA-seq and ChIP-Seq data have been deposited in GEO with the following accession numbers: bulk RNA-seq (GSE330243), epithelial cell RNA-seq (GSE330245), ChIP-seq (GSE330249). All other data are available to academic researchers upon reasonable request.

## Competing interests

The authors have no conflicts of interest to declare.

FXR is responsible for the overall content of the manuscript.

## MATERIALS AND METHODS

### Mouse strains

Suppl. Table 13 describes the mouse alleles/strains used in this study. Mice were maintained in a mixed, predominantly C57BL/6, background. Experiments were performed using mice of both sexes; age was as indicated in the text. Breedings were set in heterozygosity and littermates were used as controls. Mice of all genotypes were co-housed. In all experiments, control mice harbouring wild-type versions of the alleles shown in bold were included (short name in parenthesis). All animal procedures were approved by local and regional ethics committees [(Institutional Animal Care and Use Committee and Ethics Committee for Research and Animal Welfare, Instituto de Salud Carlos III) (CBA 09_2015_v2) and Comunidad Autónoma de Madrid (ES280790000186)] and performed according to the European Union guidelines.

### Induction of recombination in i-KG4C mice

To induce recombination, 3 doses of 4-hydroxytamoxifen (TMX) were administered via gavage to 8–10 week old mice: days 1 and 3 (10 mg diluted in 100μl of a solution containing 10% EtOH, 90% saline); day 5 (5 mg in 50 μl of the same solution). After the last dose of TMX, mice were free of treatment at least for 1 week before proceeding with further experimental manipulation.

### Induction of acute pancreatitis

Two acute pancreatitis protocols were used, mild and severe. To induce a mild acute pancreatitis, seven hourly intraperitoneal injections of caerulein (50μg/kg) were administered; saline-treated mice were used as controls. The severe acute pancreatitis protocol consisted of seven hourly intraperitoneal injections of 80 μg/kg of caerulein or PBS for two consecutive days. Mice were sacrificed by cervical dislocation at defined time points. The pancreas was processed rapidly: one piece was flash frozen and stored at -80 °C for RNA extraction and the rest was fixed for histological analysis. A minimum of 3 mice/group were included for each timepoint.

### Induction of chronic pancreatitis

Chronic pancreatitis was induced in i-KG4C mice, which had previously undergone recombination at 8-10 weeks (see induction of recombination section). When i-KG4C mice reached 10-12 weeks old, they were injected daily with one dose of caerulein (0.25mg/kg in 100μl of saline) to induce pancreatitis for 6 months. Upon pancreatitis completion, mice were aged for 3 additional months and then sacrificed for histological analysis.

### Intrapancreatic adenoviral administration

Mice (10-week-old, n=5/group) were anesthetized with isoflurane and a small incision was performed in the upper left quadrant abdominal wall to expose the pancreas. An suspension of adenovirus (1×10^9^ plaque-forming units [pfu] in 50μl saline) encoding either GFP (Ad-EGFP) or IL17A (Ad-IL17) (Schwarzenberger et al., 1998) was injected directly into multiple sites of the pancreas.

### Acinar cell isolation and culture

Acinar cell fractions (C57BL/6 mice, age 8-12 weeks) were obtained as described (Means et al., 2005). Cells were cultured in suspension in RPMI 1640 glutamax medium supplemented with 10% fetal bovine serum, penicillin, streptomycin, geneticin sulfate, and soybean trypsin inhibitor. Isolated acinar cells were cultured as described and, after 24h, they were processed for RNA extraction using Mammalian Total RNA Miniprep Kit (Sigma) according to manufacturer’s indications.

### Pancreatic epithelial cell isolation for RNA-Seq

Samples were processed as previously described (Alonso-Curbelo et al., 2021) except that Collagenase D (11088858001, Sigma-Aldrich) was used instead of Collagenase V (C9263, Sigma-Aldrich).

### *In vitro* ADM assays

Isolated acinar cells were resuspended in complete medium supplemented with 1% FBS and growth factor-reduced matrigel (1:1) and the following factors: were added at the indicated concentrations: EGF (50ng/ml), IL-17 (50ng/ml), IL-6 (50ng/ml), IL-1b (50ng/ml), IL-2 (50ng/ml), IL-4 (50ng/ml), IL-10 (2ng/ml), REG3B (500nM), Ccl5 (50ng/ml), Cxcl12 (50ng/ml). Medium was changed after 48h (day 2) and the number of cysts/well was quantified at day 4 visually under the microscope.

### Secretome analysis

Primary acinar cells were cultured for 48 h the conditioned media was collected and concentrated 10-fold using Amicon Ultra 0.5 mL Centrifugal Filters, 10 kDa MWCO (UFC501024, Merck). Concentrated secretome samples were solubilized in 5% SDS, 100 mM triethylammonium bicarbonate (TEAB) pH 8.0. and digested using the Protifi™ S-Trap™ Mini Spin Column Digestion Protocol. Peptides were analyzed by LC-MS/MS using a nanoLC-Ultra 1D+ system and an Impact mass spectrometer (Bruker). Peptides were separated on a trap column and an analytical column using a 90 min gradient. The mass spectrometer operated in data-dependent mode, switching between MS and MS/MS scans. An active exclusion of 30 sec was used. Peptides were isolated and fragmented using collision induced dissociation (CID) with a collision energy of 23-56 eV as function of the m/z value. Raw files were processed with MaxQuant (v 1.6.0.16) using the standard settings against a mouse protein database (UniProtKB/Swiss-Prot and TrEMBL, August 2015, 43,539 sequences) supplemented with contaminants. Minimal peptide length was set to 7 amino acids and a maximum of two tryptic missed-cleavages were allowed. Results were filtered at 0.01 FDR (peptide and protein level).

### Tissue processing and histopathological analysis

Mouse pancreata were fixed in 4% PBS-buffered formaldehyde overnight, embedded in paraffin, and serially sectioned (3μm). Sections from ≥3 representative levels of the tissue block (>200μm apart) were deparaffinized, H-E-stained and analyzed. Standard Alcian blue/Periodic acid-Schiff/Hematoxylin stain was performed in an automated platform (ArtisanLink Pro, Dako) according to manufacturer’s indications. Images were acquired with a Nikon Eclipse Ti microscope and managed with NIS-Elements BR3.2 software. Scoring was performed by a board-certified pathologist (IE, MI). ADM and PanIN low- (PanIN 1-2) and high-grade (PanIN 3) were identified according to established criteria (Hruban et al., 2006). AFL have been described previously (Aichler et al., 2012); briefly, they are ductular lesions occurring in ADM areas with cytological atypia (enlarged and mostly hypochromatic nuclei, visible nucleoli) displaying a characteristic “whirling” fibrosis rich in αSMA-positive myofibroblasts (Yavas et al., 2026). Remodelling refers to the areas (expressed as percentage of the whole tissue) characterized by loss of acinar tissue, ADM, fibrosis, inflammation and, sometimes, AFL. PanIN may be present, as well as cancer (as stated in the table). Tissue remodelling was defined as the occurrence of ADM, AFLs, or PanINs. The involvement by ADM/PanIN or by lipomatosis (and its degree) was semi-quantified by visual inspection.

### Immunohistochemistry

Sections of FFPE tissues were deparaffinized, rehydrated and boiled in 10mM sodium citrate buffer (pH 6.0) for 10 min for antigen retrieval. Next, sections were incubated for 30 min with 3% H_2_O_2_ in methanol to block endogenous peroxidase, washed with distilled water, and incubated for 30 min with 2% BSA in PBS at room temperature. Afterwards, sections were incubated with primary antibodies for 1h at room temperature or overnight at 4 °C, washed in PBS x3, and incubated for 45min with EnVision+ HRP labeled secondary anti-rabbit antibodies (Dako). After washing, reactions were revealed with DAB, and sections were counterstained with hematoxylin, and mounted. Suppl. Table 14 lists the primary antibodies used. Images were acquired with a Nikon Eclipse Ti microscope and managed with NIS-Elements BR3.2 software. Quantification of MCM4+ and Ki-67+ cells in control and *Gata4*^P-/-^ pancreata was performed by counting the proportion of positive cells in 5 random fields in the pancreas of ≥4 mice/genotype.

### Immunofluorescence

Tissue sections were processed as described above, skipping the H_2_O_2_ blocking step. After incubation with the primary antibody, sections were washed with PBS and incubated with secondary antibodies at 1:200 dilution (Suppl. Table 14). After washing with PBS, DAPI (0.5μg/ml in dH_2_O) was added and sections were mounted using ProLong® Gold Antifade Reagent (Life Technologies). Images were acquired using a confocal microscope (TCS SP5, Leica).

### RNA isolation

Samples were collected immediately after sacrificing the animal and flash-frozen indry ice. Frozen tissue was ground in a mortar and then homogenized using T10 basic ULTRA-TURRAX homogenizer (IKA) in GTC buffer (4M Guanidine thiocyanate, 0.1M Tris-HCl (pH 7.5), 1% 2-mercaptoethanol, prepared in DEPC-treated water). Total RNA was extracted using the phenol-chloroform method and treated with DNase (DNA-free™ DNase Treatment & Removal Reagents, Ambion). RNA was extracted from isolated epithelial cells lysed in TRIzol LS following manufacturer’s instructions with the following amendments: 1) Samples were incubated at -20 °C overnight after isopropanol addition, 2) RNA pellet was dissolved in 15μL of nuclease-free water. RNA quality was assessed by determining RNA Integrity Number (RIN) using Agilent 2100 Bioanalyzer.

### RT-qPCR

RNA was retro-transcribed with TaqMan® Reverse Transcription Reagents kit using Oligo-dT primers (ThermoFisher Scientific). Quantitative PCR was performed using SYBR® Green PCR Master Mix (ThermoFisher Scientific) in Applied Biosystems™ 7500 Fast or QuantStudio 6 Flex Real-Time PCR Systems (ThermoFisher Scientific). Gene expression was summarized as fold-change in comparison to control samples and normalized to *Hprt* levels. Oligonucleotide primers used are listed in Suppl. Table 15.

### ChIP-qPCR

Abundance of target DNA in material subject to ChIP was assessed by qPCR using the following primers (Suppl. Table 15).

### RNA-seq analysis

RNA samples with RIN>8.0 were used for RNA-seq. For Control and Gata4P-/- mice, 3 samples from each cohort were mixed to generate 3 pooled samples. For KC and KG4C mice, 3 individual samples from each experimental cohort were used. Total RNA (1 µg) containing ERCC ExFold RNA Spike-In Mixes (Thermo) was used for preparation of libraries. PolyA+ fraction was purified and randomly fragmented, converted to double stranded cDNA, and processed through subsequent enzymatic treatments of end-repair, dA-tailing, and ligation to adapters as in Illumina’s “TruSeq Stranded mRNA Sample Preparation Part # 15031047 Rev. D” kit (Illumina). Adapter-ligated library was completed by PCR with Illumina PE primers (8 cycles). The resulting purified cDNA library was applied to an Illumina flow cell for cluster generation and sequenced on Illumina HiSeq 2000 System (Illumina) according to manufacturer’s protocols (single-end read, 50 bases). Image analysis, per-cycle basecalling and quality score assignment was performed with Illumina Real Time Analysis software (Illumina). Conversion of Illumina BCL files to BAM format was performed with the Illumina2bam tool (Broad Institute of MIT and Harvard). Total number of reads ranged from 33 to 46 million reads/sample. For isolated epithelial cells RNA, libraries were prepared from total RNA, using TruSeq Stranded mRNA HT kit (llumina, Cat. N. 20020595). Sequencing was performed by Genewiz (Azenta, USA). An average of 28 million paired-end reads was generated per sample. The NExtPresso pipeline v1.9.2.5 (Graña et al., 2018) was used to process RNA-seq data. Briefly, base calling and cross contamination QC analysis were performed by FastQC and fastqscreen softwares, respectively. Reads were aligned to mm10 mouse genome using bowtie and tophat aligners. Gene count matrices were generated using htseq-count software. Normalization by median of ratios and differential gene expression analysis was performed using DEseq2 R package.

### Bioinformatics analysis of transcriptomic data

BAM files were processed and aligned to GRCm38 (mm10) mouse genome assembly using NextPresso v1.9.2.5 pipeline (Graña et al., 2018). Expression levels of features were quantified using HTseq-count as “raw counts” (Anders et al., 2015). Normalized counts and differential gene expression were estimated in R software environment (version 4.5.1) [R Core Team, 2023; https://www.R-project.org/] using DEseq2 package for R (version 3.21) (Love et al., 2014) with default model settings. Nominal p-values were adjusted for multiple testing using the false-discovery rate (FDR) correction of Benjamini and Hochberg. At least 2-fold differences between the experimental groups with FDR-adjusted p-values < 0.1 were considered biologically relevant and statistically significant. Single-cell RNA-seq data from Aney et al., 2024 were provided by the authors as a processed and annotated dataset and analyzed using Seurat package (version 5.2.0) (Hao et al., 2024). Gene signatures for specific cell populations were compiled using up to 100 most specific markers identified via scRNA-seq (Suppl. Table 7). Signatures for clusters of differentially expressed genes (DEGs) were compiled using up to 100 genes identified via partial least square discriminant analysis in mdatools package (version 0.14.2) (Kucheryavskiy, 2020). (Suppl. Table 10). Pathway enrichment analysis was performed in GSEA software using the full gene matrix (Mootha et al., 2003; Subramanian et al., 2005), with HALLMARK and REACTOME gene set collections from MSigDB (Castanza et al., 2023), an acinar-specific gene signature (Masui et al., 2010), med_rank_KRAS_ERK signatures (Klomp et al., 2024), or custom gene signatures described above.

### Chromatin immunoprecipitation (ChIP)

Pancreas from 10-week old mice were minced in cold PBS supplemented with 2x protease inhibitors, washed twice in PBS supplemented with protease inhibitors and cross-linked with 1% formaldehyde for 20 min at room temperature. Cross-linked pancreata were harvested in 0.5% SDS lysis buffer (1mL/pancreas), passed through a tight douncer, and sonicated in a Covaris instrument (shearing time 40 min, 20% duty cycle, intensity 10, 200 cycles per burst and 30sec per cycle). ChIP was then performed using anti-Gata4 antibodies (R&D systems, MAB2606). DNA was quantitated by fluorometry and the electrophoretic fraction of 100-200bp was processed through subsequent enzymatic treatments for fragmentation (5 min), end-repair, dA-tailing, and ligation to adapters with “NEBNext Ultra II FS DNA Library Prep Kit for Illumina” (New England BioLabs). Adapter-ligated library was completed by limited-cycle PCR and extracted with a double-sided SPRI size selection. Resulting average fragment size is 300 bp from which 120 bp correspond to adaptor sequences. The resulting purified DNA library was applied to an Illumina flow cell for cluster generation (TruSeq cluster generation kit v4) and sequenced on Illumina HiSeq 2500 with v4 Chemistry following manufacturer’s recommendations (single read, 50 bases). Image analysis, per-cycle basecalling and quality score assignment was performed with Illumina HiSeq Control Software. Conversion of BCL files to FASTQ format was performed with the bcl2fastq Software (Illumina).

### ChIP-seq analyses

ChIPseq analysis was carried out using Rubioseq pipeline (PMID: 27886717). Reads were mapped to reference genome (mm10) using BWA aligner and duplicates marked with Picard software. Peak calling was performed by MACS2 with the following arguments ‘--nomodel --extsize 200 --gsize mm’. Peak quality checking was done using Phantompeaks. BigWig files were generated by deeptools for peak visualization in the IGV browser. Consistent peak sets between replicates were established with mergePeaks and those regions were annotated with annotatePeaks.pl, both commands from Hypergeometric Optimization of Motif EnRichment (HOMER) software (Heinz et al., 2010). Motif analysis was performed with findMotifsGenome.pl -size given command from HOMER.

### Statistical analyses

For quantitative variables, data are provided as mean ± SD. Comparisons were performed using two-sided Mann-Whitney test, where n<5 and data did not follow a normal distribution, and two-tailed Student’s t test when n≤5. For qualitative variables, comparisons were performed using Chi-square test; when n<5, Fisher’s exact test was used. P<0.05 was considered significant. Statistical analyses were performed using GraphPad PRISM software package version 6.10 (GraphPad Software, USA). Plots were generated using GraphPad PRISM version 6.10, Morpheus software (https://software.broadinstitute.org/morpheus), and R software environment.

## Supplementary Table index

**Suppl. Table 1**. Differential gene expression between bulk pancreata of *Gata4^P-/-^* vs control mice and *Gata6^P-/-^* vs control mice.

**Suppl. Table 2**. Gene signature enrichment analysis (GSEA) performed for bulk pancreata of *Gata4^P-/-^* vs control mice using HALLMARK and REACTOME geneset libraries and the signature of pancreatic secreted enzymes reported in Masui et al., 2010.

**Suppl. Table 3**. ChIP-seq analysis of GATA4 and GATA6 in bulk pancreata of control mice.

**Suppl. Table 4**. Differential gene expression between bulk pancreata of KC and control mice. Differential gene expression between bulk pancreata of KG4C and KC mice.

**Suppl. Table 5**. Clusters of DEGs identified in comparisons of bulk pancreata of control, KC, and KG4C mice.

**Suppl. Table 6**. Gene signature enrichment analysis (GSEA) performed for bulk pancreata of KC vs control mice (and KG4C vs KC mice) using HALLMARK geneset library and MED_RANK_KRAS_ERK signatures reported in Klomp et al., 2024.

**Suppl. Table 7**. Gene signature enrichment analysis (GSEA) of signatures specific for major cell populations present in pancreatic tissue. Cell type-specific marker genes extracted from scRNA-seq data published by Aney et al., 2024.

**Suppl. Table 8**. Gene signature enrichment analysis (GSEA) performed for bulk pancreata of KC vs control mice (and KG4C vs KC mice)using cell type-specific genesets extracted fom Aney et al., 2024.

**Suppl. Table 9**. Differential gene expression in pancreatic epithelial cells isolated from control, KC, and KG4C mice. Cluster numbers and genes of interest are derived from transcriptomic analysis of bulk pancreatic tissue. Genes displaying expression changes similar to those identified in bulk pancreata are highlighted in green.

**Suppl. Table 10**. Representative GSEA gene signatures for DEG clusters identified in bulk pancreata RNA-seq analysis. Up to 100 genes that contributed most to the expression pattern of each cluster were identified by Partial Least Squares Discriminant Analysis.

**Suppl. Table 11**. Gene signature enrichment analysis (GSEA) performed for bulk pancreata of KC vs control mice using representative gene signatures for DEG clusters identified by Partial Least Squares Discriminant Analysis.

**Suppl. Table 12**. Differential protein abundance in conditioned medium from KG4C and KC primary acinar cells.

**Suppl. Table 13**. Mouse alleles used in this study.

**Suppl. Table 14**. Antibodies used in this study.

**Suppl. Table 15**. Oligonucleotide primers used for RT-qPCR and ChIP-qPCR.

## Notes

### Competing Interest Statement

The authors have declared no competing interest.

